# Feeding induces c-Fos in hepatocytes contributing to hepatocellular carcinoma in obesity

**DOI:** 10.1101/2025.05.20.654880

**Authors:** Ao Li, Eduardo H. Gilglioni, Wadsen St-Pierre-Wijckmans, Leila Hosseinzadeh, Christelle Veyrat-Durebex, Sumeet P. Singh, Roberto Coppari, Latifa Bakiri, Esteban N. Gurzov

## Abstract

The transcription factor c-Fos plays an important role in hepatic metabolism; however, its role in metabolic dysfunction-associated steatotic liver disease (MASLD) and hepatocellular carcinoma (HCC) is unclear. Here, we show that hepatic c-Fos is induced by insulin after feeding and suppressed by glucagon during fasting in chow-fed mice. In lean mice, adenovirus-mediated c-Fos ectopic expression in the liver is sufficient to cause insulin resistance. In diet-induced obesity or after ectopic expression in hepatocytes, c-Fos promotes MASLD progression by altering PPAR signaling and fatty acid metabolism pathways. Mechanistically, c-Fos drives glycolysis, stress-associated MAPK, and insulin-related PI3K-Akt signaling, exacerbating metabolic dysregulation. In HCC, c-Fos expression correlates with PI3K-Akt, MAPK, and calcium signaling pathways activation. Moreover, c-Fos siRNA knockdown in human liver cancer cells reduces proliferation and increases apoptosis under lipotoxic or ER stress conditions. These findings identify c-Fos as a critical mediator of liver steatosis progression, linking hepatocyte signaling and metabolic reprogramming to liver dysfunction and tumorigenesis.

## Introduction

Obesity affects over 1 billion people worldwide and is a major contributor to metabolic-associated steatotic liver disease (MASLD), which impacts approximately 38% of the global population [1]. MASLD can progress to severe liver diseases as metabolic-associated steatohepatitis (MASH), cirrhosis, and hepatocellular carcinoma (HCC) [2]. The liver, as the central hub for metabolic homeostasis, is particularly affected during obesity [3]. In recent years, the obesity epidemic has been implicated in up to 40% of the increasing incidence of HCC in developed nations [4].

The liver functions by integrating hormonal, nutritional, and neuronal cues to regulate hepatocyte metabolic activity. During fasting, it prioritizes fatty acid oxidation and maintains glucose levels through glycogenolysis and gluconeogenesis, while carbohydrate intake stimulates hepatic glucose uptake, glycogen synthesis, glycolysis, and lipogenesis [5, 6]. These metabolic shifts are tightly controlled by insulin, glucagon, and other hormonal cues, adjusting dynamically to nutrient availability and the body’s energy needs [6]. However, chronic overnutrition and obesity disrupt this finely tuned regulatory network, driving hepatic steatosis, inflammation, and metabolic reprogramming that are key contributors to liver disease progression [2, 7].

Numerous transcription factors regulate hepatocyte metabolism and contribute to the progression of MASLD, MASH, and HCC. Among them, the role of the dimeric activator protein 1 (AP-1) transcription factor family in cell growth, differentiation, apoptosis, and stress responses is well documented [8, 9]. c-Fos, the first discovered AP-1 member [10], plays a particularly prominent role, orchestrating adaptive metabolic and signaling responses across different cell types. c-Fos expression is a marker of neuronal activation in circuits associated with appetite regulation, including pro-opiomelanocortin (POMC) neurons [11] and corticostriatal circuits [12], both of which influence feeding behavior. In β-cells, c-Fos modulates insulin secretion [13]. In adipocytes, c-Fos facilitates adipogenesis by enhancing the expression of key transcription factors such as Peroxisome Proliferator Activated Receptor (PPAR) γ [14]. In hepatocytes, c-Fos activates PPARγ and contributes to the development of MASH and HCC in response to palmitate-rich high-fat diets [15–17]. Transcriptomic studies further highlight c-Fos as a key protein regulating multiple signaling pathways involved in MASLD [18]. However, the role of c-Fos in hepatocyte glucose and lipid metabolism and its contribution to liver dysfunction and disease progression in obesity are still not completely understood.

Here, we studied hepatic c-Fos expression in chow diet-fed and obesogenic diet-fed mice under fasting and feeding conditions. Insulin stimulates c-Fos expression in hepatocytes, while obesogenic diet components, such as saturated fatty acids and fructose, enhance its expression. Increased c-Fos alters the expression of genes regulating glucose and lipid metabolism, leading to metabolic changes associated with MASLD and HCC progression. Our findings establish c-Fos as a key regulator of hepatic energy metabolism and underscore its role in metabolic liver diseases in obesity.

## Results

### Hepatic c-Fos is induced by insulin during feeding

c-Fos is rapidly induced in response to stimuli and undergoes gradual degradation [19]. To explore its fluctuations during fasting and feeding, we first analyzed a publicly available RNA sequencing dataset (GSE200811) derived from the livers of humanized male chimeric mice sampled every 4 hours over the circadian cycle (Zeitgeber Time 0, ZT0: lights on; ZT12: lights off) [20]. *FOS* mRNA levels remained low and stable throughout the light phase (resting/fasting period) but increased at the onset of the dark phase, when mice are active and typically begin feeding (**Fig. S1A**). To directly investigate feeding-induced c-Fos expression, 8-week-old C57BL/6N lean mice were randomly assigned to one of two experimental conditions: a “24-hour-Fasted” protocol (Fasted group, ZT0 to ZT24) or a “12-hour-Fasted (ZT0-12, lights on) & 12-hour-Fed” protocol (Fed group, ZT12-24, lights off) (**Fig. 1A**). At end point (ZT24), the fed group exhibited higher body weight, glycemia, and insulinemia when compared to the fasted group (**Fig. 1B**). Liver weight, liver-to-body weight ratio, liver lean mass, and liver total water content were also significantly higher in the fed group. No significant differences were observed in liver fat mass. Histological analyses showed larger hepatocytes in the fed group (**Fig. S1B**), with increased glycogen accumulation (**Fig. 1C**). Increased c-Fos protein was detected by immunoblotting in the fed group, as well as higher protein expression of the lipogenesis-associated markers PPARγ and PLIN2 (**Fig. 1D**). In line with increased insulinemia, phosphorylation of the insulin receptor (IR), AKT at Ser473, and ERK1/2 was increased in the fed group. At the mRNA level, β-oxidation-related *Pparα* and its downstream gene *Cpt1α*, as well as antioxidant responsive *Nfe2l2* were decreased in the fed group, while lipogenesis-related *Pparγ* isoform 2 and its target genes (*Acly*, *Acaca*, and *Fasn*) were significantly upregulated (**Fig. 1E**). The upregulation of lipogenic markers and insulin signaling pathways, along with the downregulation of fatty acid oxidation genes, suggested a shift toward anabolic lipid metabolism in the fed state, which follows the increase in c-Fos protein.

**Fig. 1.**
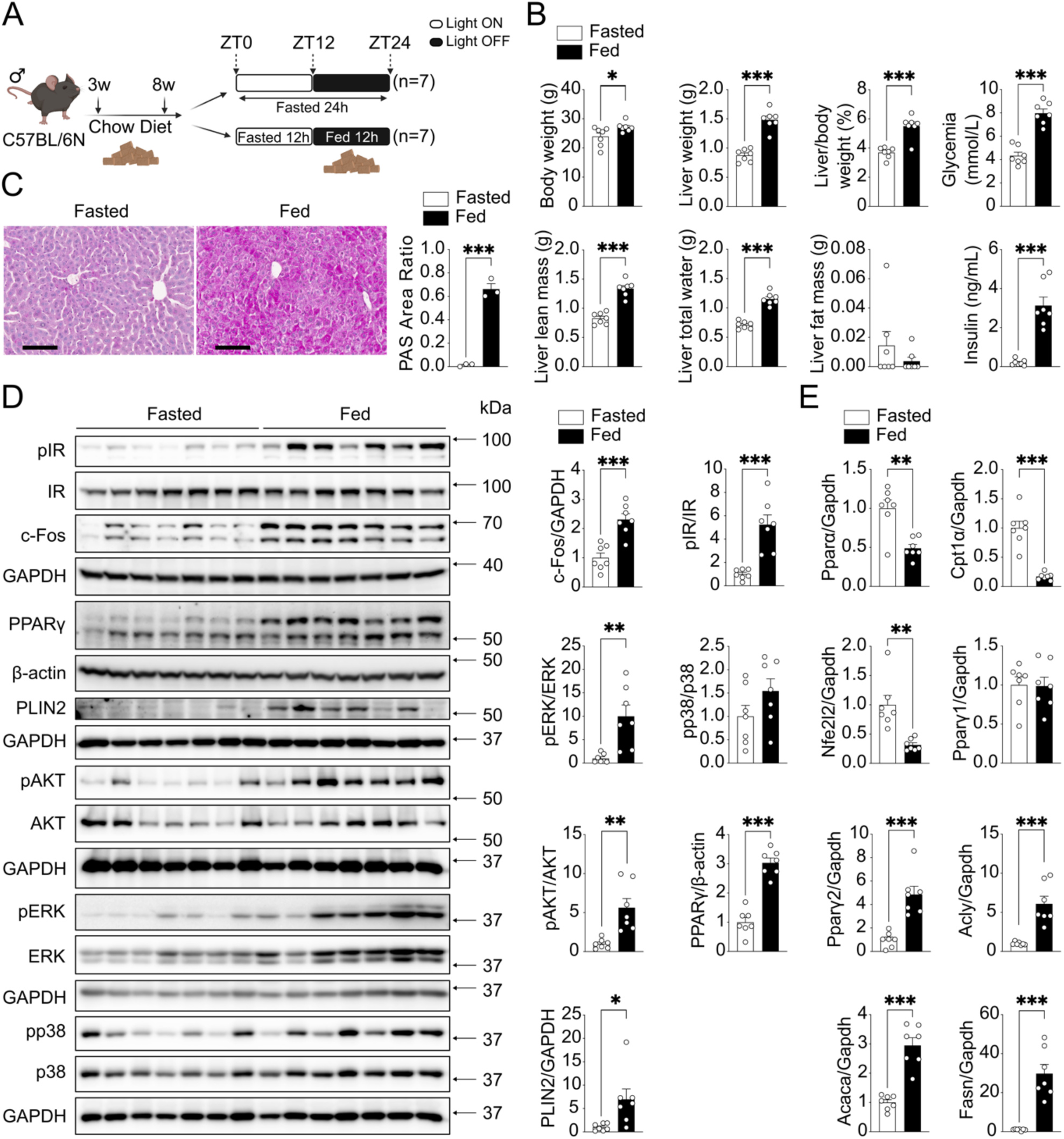
Hepatic c-Fos expression increases during feeding in chow diet-fed mice. (A) Methodological approach schematic illustrating the fasted-fed experimental protocol. Eight-week-old C57BL/6N male mice were subjected to either a 24-hour fasting protocol (Fasted group: ZT0-ZT24) or a 12-hour fasting (ZT0-ZT12) followed by 12-hour feeding (ZT12-ZT24) protocol (Fed group). (B) Metabolic differences between fasted (n=7) and fed (n=7) groups as indicated. (C) Histological analyses using PAS staining demonstrate glycogen accumulation in mouse livers (n=3). Scale bar=100μm. (D) Immunoblot analysis of fasted (n=7) and fed (n=7) mouse liver tissues showing pIR, pAKT, pERK, pp38 MAPK, c-Fos and lipogenesis markers (PPARγ and PLIN2) protein expressions. (E) RT-PCR analysis shows β-oxidation-related genes (*Pparα*, *Cpt1α*, *Nfe2l2*) and lipogenesis-related genes (*Pparγ*, *Acly*, *Acaca*, *Fasn*) in fasted/fed mouse livers (n=6-7). In (B-E), results are shown as mean ± SEM. Statistical analyses using two-tailed unpaired Student’s t-test (B-E). Statistical significance is indicated as \**p* < 0.05, \*\**p* < 0.01, \*\*\**p* < 0.001.

Next, primary mouse hepatocytes (mHep) were isolated by a two-step collagenase perfusion method. Insulin treatment of mHep resulted in a significant increase in *Fos* mRNA levels within 30 min (**Fig. 2A**). pIR protein levels were elevated within 30 min, and c-Fos protein levels significantly increased after 2h of insulin treatment (**Fig. 2B**). c-Fos is induced by insulin through the phosphorylation of ERK1/2 and p38 MAPK in several cell types [21, 22]. Thus, an ERK phosphorylation inhibitor (ERKi, SCH772984) was included with insulin in mHep cultures. pERK and c-Fos protein levels were significantly reduced in the presence of ERKi compared to insulin only after 2h of treatment (**Fig. 2C**). Immunofluorescence staining further confirmed reduced insulin-induced c-Fos expression in the presence of ERKi (**Fig. 2D**). Conversely, glucagon treatment significantly reduced pERK and c-Fos protein expression in mHep, highlighting the regulation of c-Fos by ERK under fasting conditions (**Fig. 2E**).

**Fig. 2.**
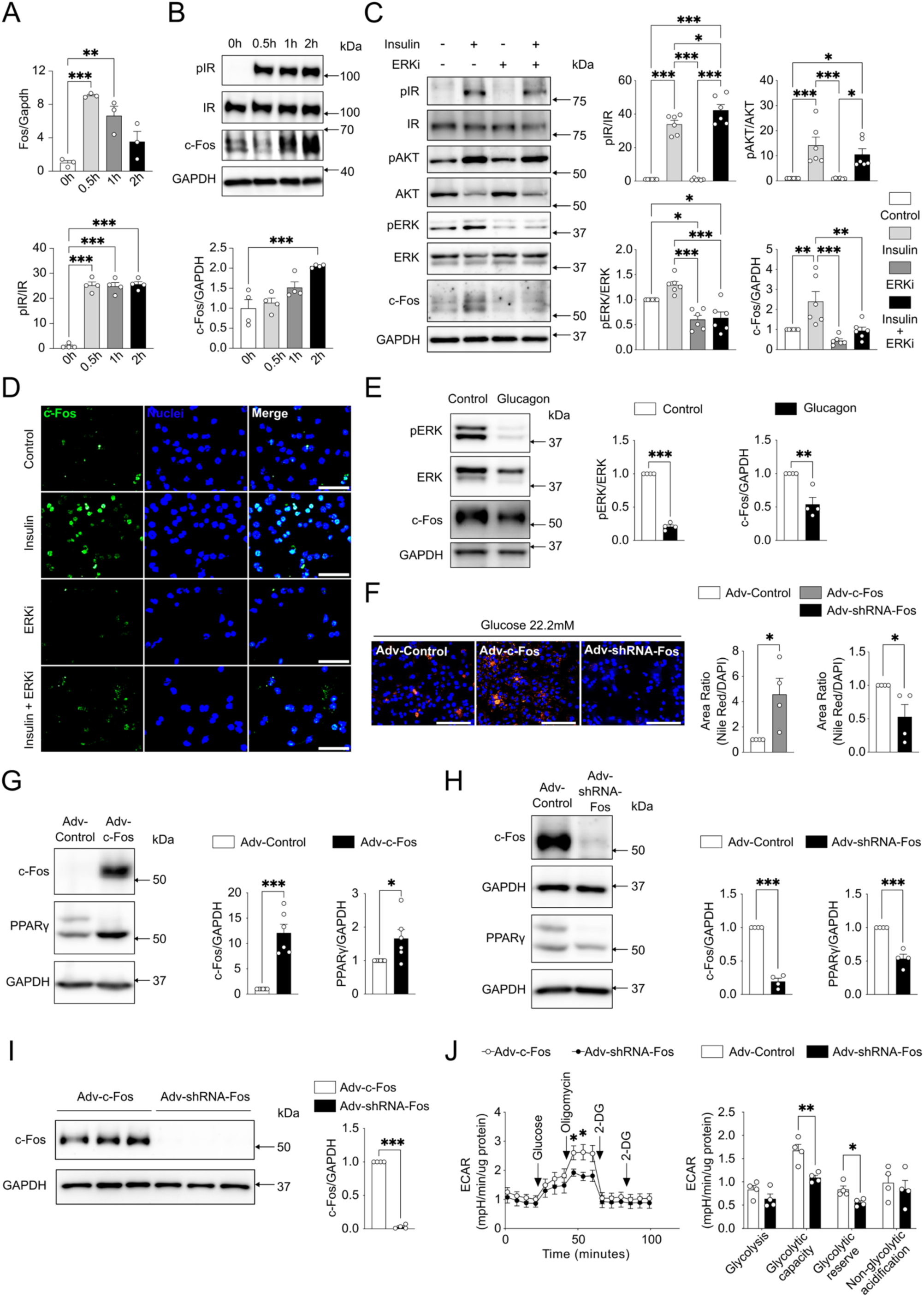
Insulin-induced c-Fos expression promotes fat accumulation in primary mouse hepatocytes. (A) RT-PCR (n=3) and (B) immunoblot analysis (n=4) indicating *Fos* mRNA and pIR, c-Fos protein expression in primary mouse hepatocytes (mHep) with 2h insulin treatment (100nM). (C) Immunoblot analysis of primary mHep (n=6) showing pIR, pAKT, pERK, and c-Fos protein expression with insulin (100nM) and an ERK inhibitor (ERKi, SCH772984, 150nM) treatment after 2h. (D) c-Fos protein expression in primary mHep with insulin (100nM) and an ERK inhibitor (ERKi, SCH772984, 150nM) treatment after 2h. Scale bar=100μm. (E) Immunoblot analysis of primary mHep (n=4) showing pERK and c-Fos protein expression with glucagon (100nM) treatment after 2h. (F) Nile Red and DAPI staining demonstrate lipid accumulation in primary mHep (n=4) with adenovirus-mediated c-Fos overexpression (Adv-c-Fos, 50MOI) or silencing (Adv-shRNA-Fos, 50MOI) in high glucose (22.2mM) culture conditions for 24h. Scale bar=200μm. (G, H) Immunoblot analysis of primary mHep showing c-Fos and PPARγ protein expression with Adv-c-Fos (50MOI, n=6) or Adv-shRNA-Fos (50MOI, n=4) and high glucose (22.2mM) treatment after 24h. (I) Immunoblot analysis of primary mHep (n=4) showing c-Fos protein expression with Adv-c-Fos (50MOI) and Adv-shRNA-Fos (50MOI). (J) Real-time measurement of the extracellular acidification rate (ECAR) in response to glycolytic modulators reveals changes in glycolytic parameters of primary mHep (n=4). Each individual value represents independent hepatocyte preparations from different mice. In (A-C, E-J), results are shown as mean ± SEM. Statistical analyses using one-way ANOVA (A-C) or two-tailed unpaired Student’s t-test (E-J). Statistical significance is indicated as \**p* < 0.05, \*\**p* < 0.01, \*\*\**p* < 0.001.

Adenoviral vectors were next used to ectopically express (Adv-c-Fos) or silence (Adv-shRNA-Fos) c-Fos in mHep (**Fig. S2A**). Glucose uptake was assessed across a range of glucose concentrations simulating different physiological states of glycemia, including hypoglycemia (1.5mM), euglycemia (5.5mM), mild hyperglycemia (11.11mM), and severe hyperglycemia (22.22mM). Glucose uptake increased with elevated glucose concentrations (**Fig. S2B**), and c-Fos expression further enhanced glucose uptake at 1.5 mM, 5.5 mM, and 11.1 mM glucose. The effect was not observed at 22.2 mM glucose, but c-Fos still increased hepatocyte lipid accumulation in this setting (**Fig. 2F**). Ectopic c-Fos expression led to increased PPARγ protein and suppressed *Pparα*, *Cpt1α*, and *Nfe2l2* transcription, while conversely, c-Fos silencing lowered PPARγ (**Fig. 2G,H; Fig. S2C**). Furthermore, hepatocyte glycolytic capacity and reserve were significantly increased upon ectopic c-Fos expression compared to c-Fos silencing (**Fig. 2I,J**). Taken together, c-Fos is suppressed by glucagon and induced by insulin through ERK signaling pathway in healthy livers and isolated primary mHep. Furthermore, c-Fos stimulates glucose uptake, glycolysis, and lipid storage in hepatocytes.

### Hepatic c-Fos silencing suppresses PPARγ, and c-Fos overexpression induces insulin resistance in chow diet-fed mice

To validate our findings *in vivo*, chow diet-fed 6-week-old C57BL/6N mice were randomly assigned to c-Fos silencing (Adv-shRNA-Fos) or control (Adv-Control) groups (**Fig. 3A**). After 2 weeks, all mice underwent 12h fasted & 12h fed protocol, and the livers were collected at end point ZT24. No significant changes were detected in body weight, blood glucose, hepatic glycogen content, or liver mass and composition (**Fig. 3B; Fig. S3A**). Liver immunoblotting analysis demonstrated efficient c-Fos silencing and lower PPARγ protein levels, consistent with the *in vitro* data (**Fig. 3C**). RNA sequencing was next performed using adenovirus-infected livers (**Fig. S3B**). Genes such as the cell cycle regulator *Sik1* and the autophagy regulator *Depp1*, exhibited significant alterations in Adv-shRNA-Fos livers compared to Adv-Control, while MAPK signaling tended to be reduced (**Fig. 3D,E**). Gene Set Enrichment Analysis (GSEA) [23, 24] of Kyoto Encyclopaedia of Genes and Genomes (KEGG) in Adv-shRNA-Fos/Adv-Control revealed several enriched pathways, including suppressed MAPK and Wnt signaling, as well as activated mitochondrial complex UCP1 in thermogenesis and oxidative phosphorylation pathways (**Fig. 3F**).

**Fig. 3.**
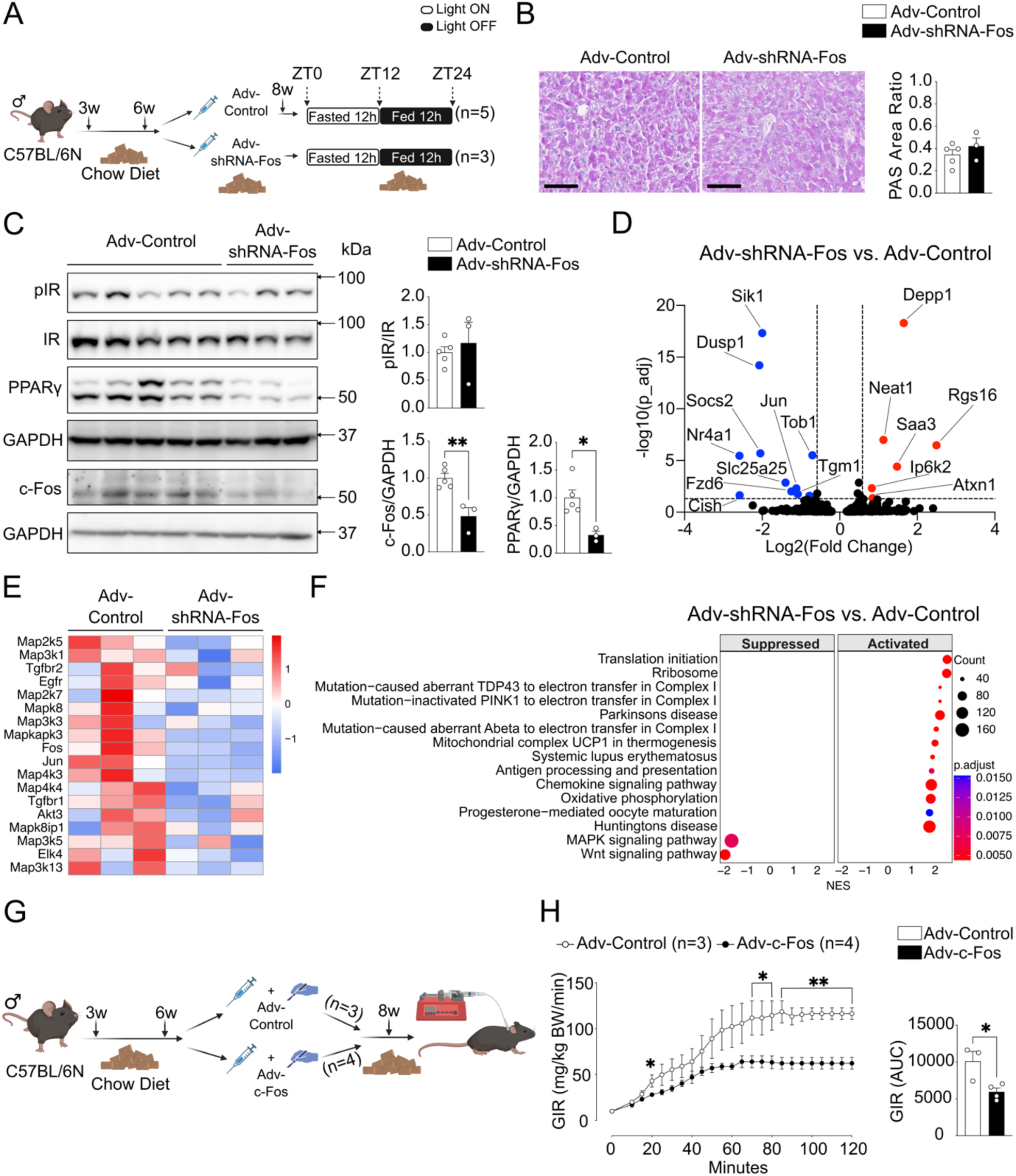
Hepatic c-Fos silencing suppresses PPARγ, and c-Fos overexpression induces insulin resistance in chow diet-fed mice. (A) Methodological approach schematic illustrating the fed experimental protocol. (B) Histological analyses using PAS staining demonstrate glycogen accumulation in Adv-Control (n=5) and Adv-shRNA-Fos (n=3) injected mouse livers. Scale bar=100μm. (C) Immunoblot analysis of Adv-Control (n=5) and Adv-shRNA-Fos (n=3) mouse livers showing pIR, c-Fos and PPARγ protein expressions. (D) RNA-Seq volcano plot displaying the quantification of transcripts between Adv-Control (n=3) and Adv-shRNA-Fos (n=3) mouse livers. (E) RNA-Seq heatmap displaying alterations in MAPK pathway-related genes. (F) RNA-Seq KEGG pathway enrichment analysis comparing Adv-shRNA-Fos vs. Adv-Control mouse livers. (G) Methodological approach schematic illustrating the hyperinsulinemic-euglycemic clamp experimental protocol. (H) Glucose infusion rate (GIR) and area under curve (AUC) between Adv-Control (n=3) and Adv-c-Fos (n=4) as indicated. In (B, C, H), results are shown as mean ± SEM. Statistical analyses using two-tailed unpaired Student’s t-test (B, C, H). Differential expression analysis using DESeq2 and pathway enrichment analysis using fGSEA with Benjamini-Hochberg FDR correction (D, F). Statistical significance is indicated as \**p* < 0.05, \*\**p* < 0.01.

Next, we overexpressed c-Fos in the liver of chow diet-fed 6-week-old C57BL/6N catheterized mice (Adv-c-Fos) for 2 weeks and subsequently subjected the mice to hyper-insulinemic/euglycemic clamps to assess insulin sensitivity (**Fig. 3G**). Circulating insulin levels were experimentally increased during clamp based on body weight (**Fig. S3C,D**). Decreased insulin sensitivity was revealed by a lower glucose infusion rate and lower glucose area under curve in clamped euglycemic Adv-c-Fos mice compared to controls (Adv-Control) (**Fig. 3H**). Collectively, these data demonstrate that, during feeding, c-Fos is a key regulator of PPARγ in the liver and that c-Fos promotes insulin resistance in chow diet-fed mice.

### Hepatic p38 MAPK and ERK activation enhance feeding-induced c-Fos expression in obese mice

Whether the induction of c-Fos by feeding in healthy, insulin-sensitive mice, on a chow diet is recapitulated in obese mice with steatosis and insulin resistance is not known. Thus, we subjected 8- week-old C57BL/6N mice to a high-fat, high-fructose, and high-cholesterol diet (HFHFHCD). After 12 weeks on HFHFHCD, obese mice were randomly assigned to fasted and fed groups, following a protocol similar to the one used for chow-fed mice (**Fig. 4A**). Obese mice in the fed group exhibited increased body weight, liver weight, liver-to-body weight ratio, liver lean mass, liver total water content, glycemia, and serum insulin levels (**Fig. 4B**). In contrast to lean mice, obese mice showed a significant increase in liver fat mass in fed compared with the fasted condition (**Fig. 4B**). Representative liver H&E and PAS staining revealed larger hepatocytes and increased glycogen accumulation in the fed group (**Fig. S4A; Fig. 4C**). No differences in pAKT levels at Ser473 were observed between fed and fasted groups, confirming insulin resistance in obese mice (**Fig. 4D**). The fed group had elevated levels of pIR, pERK1/2, c-Fos, PPARγ, and PLIN2, along with significant activation of the stress-related p38 MAPK. These observations indicate that c-Fos is induced by HFHFHCD feeding in obese mice and suggest that p38 activation may contribute, together with ERK, to increased c-Fos expression in this setting. While no differences were observed between fasted and fed groups in *Pparγ2* and *Cpt1α* transcripts, *Pparα and Pparγ1* were downregulated while *Acly*, *Acaca* and *Fasn* were significantly upregulated in the fed group (**Fig. 4E**).

**Fig. 4.**
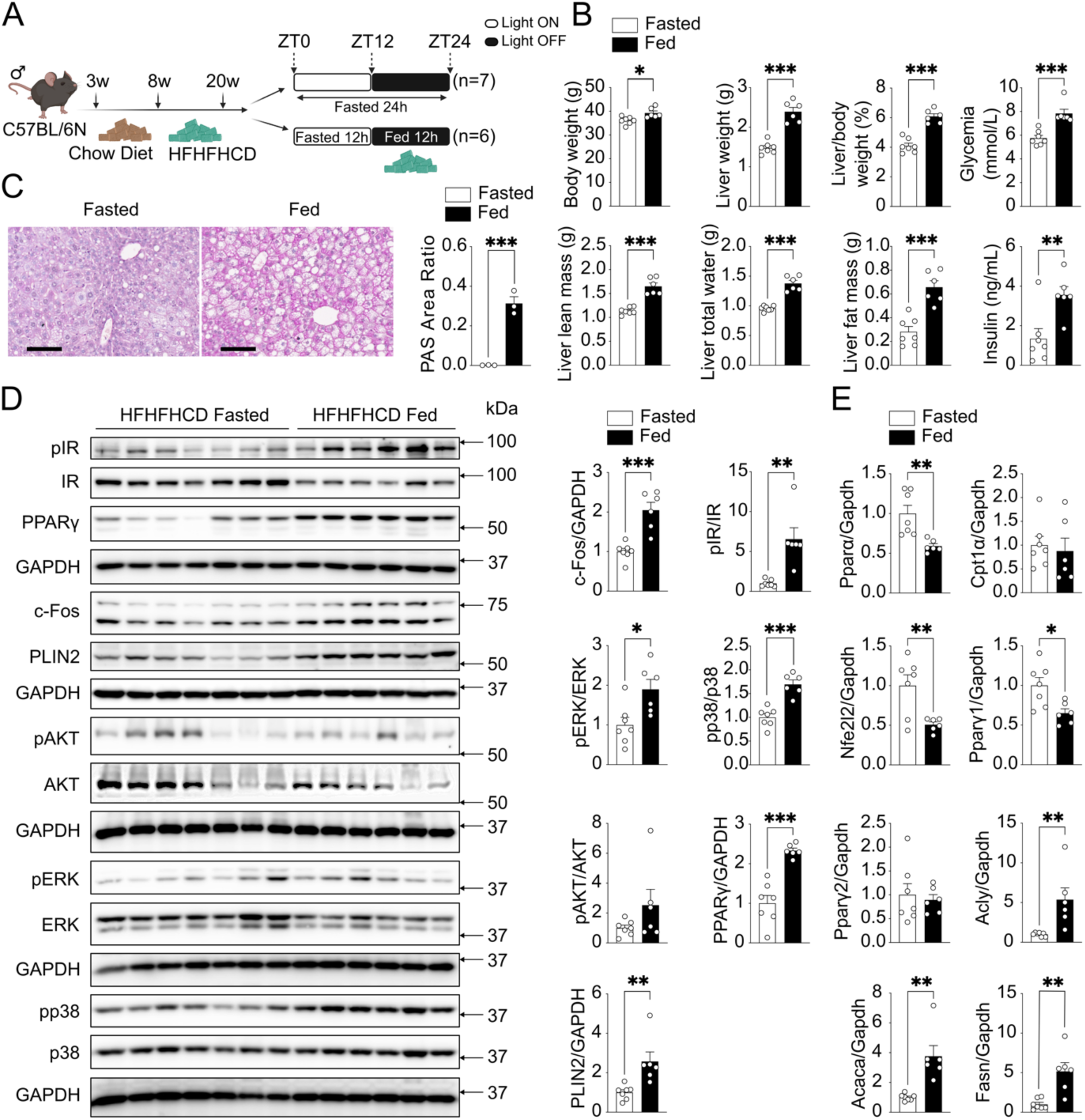
Hepatic c-Fos expression is enhanced during feeding in HFHFHCD-fed mice. (A) Methodological approach schematic illustrating the fasted-fed experimental protocol in obese mice fed with 12 weeks HFHFHCD. (B) Metabolic differences between fasted (n=7) and fed (n=6) groups as indicated. (C) Histological analyses using PAS staining demonstrate glycogen accumulation in mouse livers (n=3). Scale bar=100μm. (D) Immunoblot analysis of fasted (n=7) and fed (n=6) mouse liver tissues showing pIR, pAKT, pERK, pp38 MAPK, c-Fos and lipogenesis markers (PPARγ and PLIN2) protein expressions. (E) RT-PCR analysis shows β-oxidation-related genes (*Pparα*, *Cpt1α*, *Nfe2l2*) and lipogenesis-related genes (*Pparγ*, *Acly*, *Acaca*, *Fasn*) in fasted/fed mouse livers (n=6-7). In (B-E), results are shown as mean ± SEM. Statistical analyses using two-tailed unpaired Student’s t-test (B-E). Statistical significance is indicated as \**p* < 0.05, \*\**p* < 0.01, \*\*\**p* < 0.001.

We next compared hepatic p38 levels in mice fed with HFHFHCD for 24 weeks with mice on a chow diet (**Fig. S4B**). Immunoblot analysis showed a significant p38 activation, along with elevated c-Fos and PPARγ expression in the HFHFHCD-fed mouse livers, while ERK and AKT activation were unaffected (**Fig. 5A**). Next, isolated mHep were treated with free fatty acids, including palmitic and oleic acid, for 24h to assess the specific effects of HFHFHCD components. RT-PCR analysis revealed that palmitic but not oleic acid induced *Fos* mRNA expression (**Fig. S4C**). p38 MAPK inhibitor prevented palmitic acid-induced c-Fos protein expression in mHep (**Fig. 5B**), and this was confirmed by immunofluorescence (**Fig. 5C**). Palmitic acid treatment of mHep pre-infected with adenovirus-mediated c-Fos overexpression led to more lipid droplet accumulation and higher PPARγ expression when compared to mHep with c-Fos silencing (**Fig. 5D,E**). Interestingly, c-Fos expression was also induced by fructose but not by cholesterol, indicating that not all components of HFHFHCD increase c-Fos expression in primary mHep (**Fig. S4D**). However, fructose-induced c-Fos could not be suppressed by the p38 inhibitor suggesting independent mechanisms. Together, these results indicate that saturated free fatty acids and fructose stimulate hepatic c-Fos expression in the obese mouse liver, at least in part through p38 activation.

**Fig. 5.**
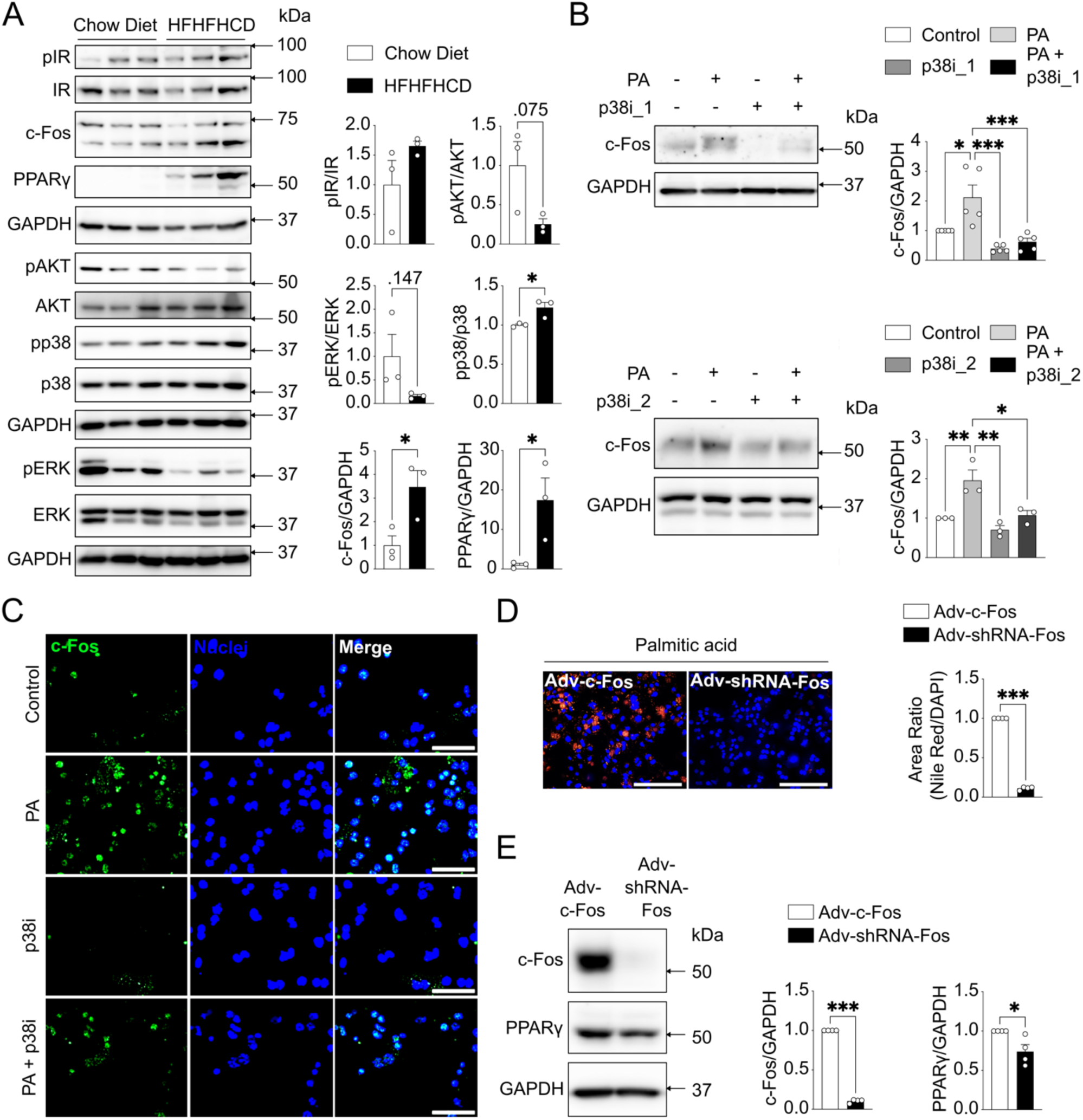
p38 MAPK and ERK activation enhance hepatic c-Fos expression in obese mice. (A) Immunoblot analysis of chow diet (n=3) and HFHFHCD fed (n=3) mouse livers showing pIR, pAKT, pERK, pp38 MAPK, c-Fos and PPARγ protein expressions. (B) Immunoblot analysis of primary mouse hepatocytes (mHep) being treated with palmitic acid (PA, 0.4mM) and p38 MAPK inhibitor 1 (Adezmapimod, p38i_1, 150nM, n=5, top) or p38 MAPK inhibitor 2 (Ergothioneine, p38i_2, 250μM, n=3, bottom) for 4h revealing c-Fos protein expression. (C) Immunofluorescence staining demonstrates c-Fos protein expression in primary mHep with PA (0.4mM) and p38i_2 (250μM) treatment after 4h. Scale bar=100μm. (D) Nile Red and DAPI staining demonstrate lipid accumulation in primary mHep (n=4) with adenovirus-mediated c-Fos overexpression (Adv-c-Fos, 50MOI) or silencing (Adv-shRNA-Fos, 50MOI) with 24h PA (0.4mM) treatment. Scale bar=200μm. (E) Immunoblot analysis of primary mHep (n=4) showing c-Fos and PPARγ protein expression after Adv-c-Fos (50MOI) and Adv-shRNA-Fos (50MOI) with 24h PA (0.4mM) treatment. In (A, B, D, E), results are shown as mean ± SEM. Statistical analyses using two-tailed unpaired Student’s t-test (A, D, E) or one-way ANOVA (B). Statistical significance is indicated as \**p* < 0.05, \*\**p* < 0.01, \*\*\**p* < 0.001.

### c-Fos overexpression in hepatocytes contributes to MASLD by modulating insulin and stress signaling pathways

We next analyzed bulk RNA sequencing datasets generated using a mouse model, where c-Fos was ectopically expressed in hepatocytes [15]. c-Fos overexpression (Fos^Hep^) was induced on chow diet at three weeks of age, and the livers were harvested for RNA sequencing analyses two months (2mo) and four months (4mo) later (**Fig. 6A**). In these mice that have a median survival of 5 months, focal liver damage is observed at the 2mo time point, whereas liver dysfunction is molecularly and histologically apparent at 4mo [15].

**Fig. 6.**
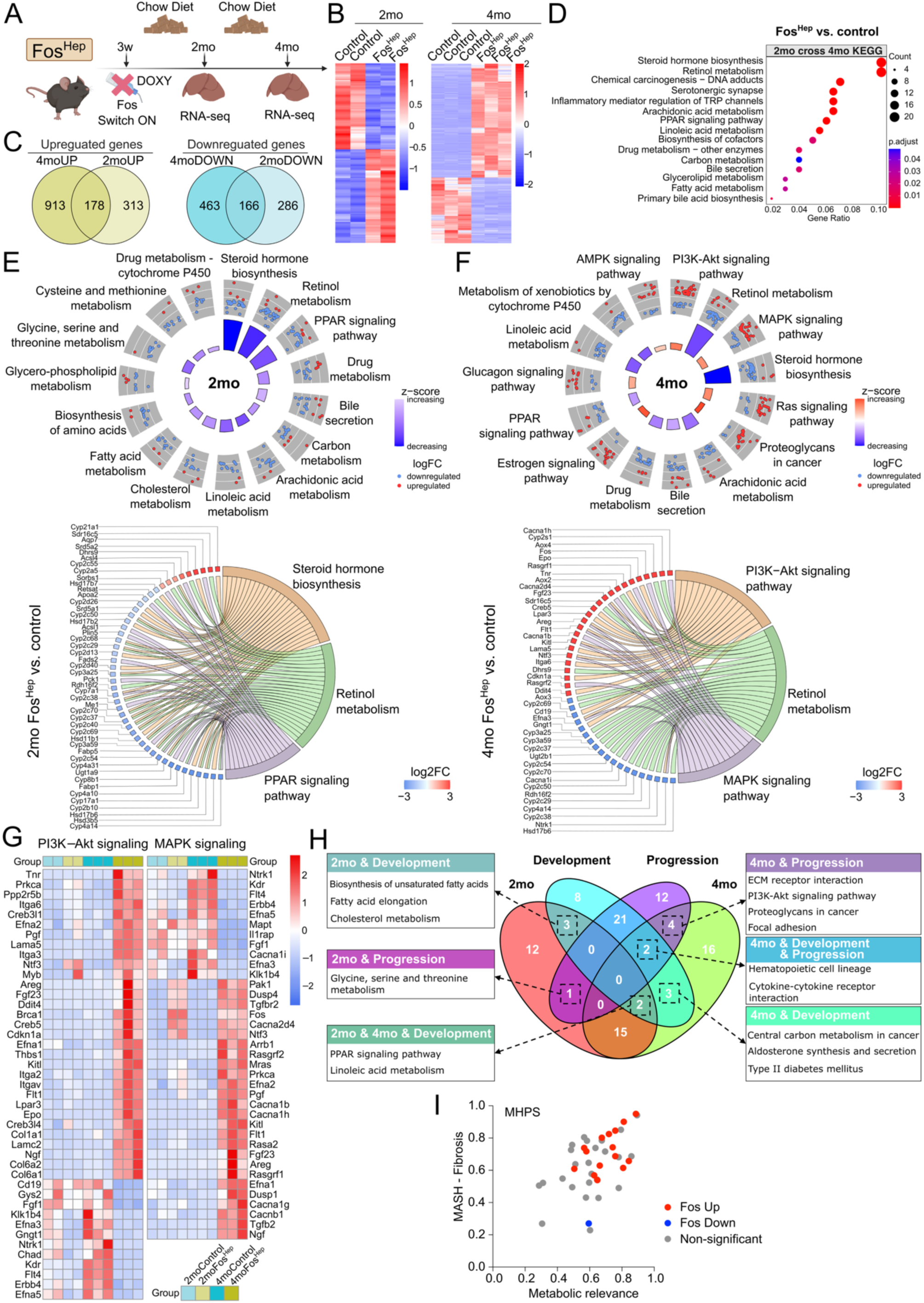
c-Fos overexpression in hepatocytes contributes to MASLD by modulating insulin and stress signaling pathways. (A) Hepatocyte-specific *Fos* (Fos^Hep^) doxycycline (DOXY)-inducible mice were subjected to RNA-Seq (GSE81079) after 2 months (2mo, n=2) and 4 months (4mo, n=3) fed with a chow diet. (B) RNA-Seq heatmaps showing significantly altered genes between control and Fos^Hep^ mice at 2mo (left) and 4mo (right). (C) RNA-Seq Venn diagrams showing the cross-up- (left) and down-regulated (right) genes between 2mo and 4mo. (D) RNA-Seq KEGG pathway enrichment analysis using the 2mo and 4mo cross-up- and down-regulated genes from (C) as indicated. (E, F) RNA-Seq circle plots showing the KEGG metabolism-related pathways in 2mo (E, top) and 4mo (F, top), comparing Fos^Hep^ vs. control, respectively. RNA-Seq chord plots showing top variable genes corresponding to the pathways in 2mo (E, bottom) and 4mo (F, bottom). Genes are ranked by log_2_FC. (G) RNA-Seq heatmap indicating gene alterations in PI3K-Akt and MAPK signaling pathways between control (2moCon, 4moCon) and Fos^Hep^ (2moFos^Hep^, 4moFos^Hep^). (H) RNA-Seq Venn diagram indicating the cross-pathways among Fos^Hep^/control (2mo and 4mo, respectively) and pathways associated with the development and progression stages of MASLD extracted from (PMCID: PMC11199145). (I) MHPS (MASLD human proximity score) graph showing the proximity between hepatic c-Fos upregulation in several rodent models and human MASLD/MASH. Data was extracted from PMCID: PMC11199145. Differential expression analysis using DESeq2 and pathway enrichment analysis using clusterProfiler with Benjamini-Hochberg (B, D, E, F). Z-SCORE quantification using GOplot (E, F). Source data is provided as a Source Data file.

Principal Component Analysis (PCA) (**Fig. S5A**) and heatmaps (**Fig. 6B**) showed significant transcriptional variations between the 2mo and 4mo Fos^Hep^ groups compared to their respective controls. A total of 178 upregulated and 166 downregulated genes were shared between the 2mo and 4mo groups (**Fig. 6C**). Over Representation Analysis (ORA) of the KEGG revealed commonly altered pathways in 2mo and 4mo Fos^Hep^ livers, including PPAR signaling, bile secretion, fatty acid metabolism, and primary bile acid biosynthesis (**Fig. 6D**). Within the PPAR signaling pathway, fold changes in β-oxidation-related *Ppara* and *Ppard*, as well as lipogenesis-related *Pparg* (**Fig. S5B**). ORA with z-score [25] and GSEA were next applied to evaluate KEGG pathways in the 2mo and 4mo groups separately (**Fig. 6E**,**F****; Fig. S5C**,**D**). Pathways such as steroid hormone biosynthesis, retinol metabolism, PPAR signaling, and bile secretion were downregulated in both Fos^Hep^ groups (**Fig. 6E**,**F**). Genes of the MAPK signaling pathway were upregulated in both Fos^Hep^ groups, with more pronounced changes observed at 4mo, as identified by both GSEA and ORA with z-score (**Fig. 6F**; **Fig. S5C**,**D**). Additionally, ectopic c-Fos expression was associated with upregulation of genes in PI3K-Akt signaling, Ras signaling, proteoglycans in cancer, estrogen signaling, and glucagon signaling (**Fig. 6F**). PI3K-Akt and MAPK pathways enrichment in Fos^Hep^ livers were highlighted with increased expression of the respective target gene (**Fig. 6G**). Given the wide range of pathways altered by c-Fos, we compared the 2mo and 4mo ORA results with pathways implicated in the development and progression stages of MASLD, as detailed in [18]. Overlapping pathways, such as PPAR and PI3K-Akt signaling, associated with development and progression stages of MASLD and Fos^Hep^ are indicated in Venn diagram (**Fig. 6H**). Additionally, a strong association between hepatic c-Fos upregulation and liver dysfunction in MASLD/MASH is apparent when mining metabolic relevance and MASH-Fibrosis scores, replotted from [18] (**Fig. 6I**).

Overall, these results suggest that increased hepatic c-Fos expression promotes MASLD through multiple signaling pathways in the liver, including insulin-associated PI3K-Akt and stress-associated MAPK signaling.

### c-Fos contributes to dedifferentiation and proliferation in HCC cells

Hepatocyte metabolic reprogramming and inflammation driven by c-Fos have been linked to liver disease and to the initial steps of HCC in mouse models [15]. To examine c-Fos expression in established HCC, we administered diethyl nitrosamine (DEN) to 2-week-old male mice, which were subsequently fed a 37-week chow diet (**Fig. S6A**) or a 25-week HFHFHCD (**Fig. S6C**), respectively. Hepatic c-Fos expression positively correlated with insulin receptor phosphorylation in both chow and HFHFHCD-fed models, and with AKT phosphorylation only in chow-diet-fed mice (**Fig. S6B,D**).

During neoplastic transformation to HCC, hepatocytes dedifferentiate and acquire a proliferative phenotype resembling progenitor stem cells. Previous studies demonstrated that c-Fos gene inactivation facilitates hepatocyte differentiation in both mouse and human liver organoids [26]. In a human hepatocyte-like cell (HLC) differentiation model (**Fig. 7A**), we observed a progressive decrease in c-Fos expression, while albumin levels significantly increased at the hepatocyte maturation stage (**Fig. 7B**), consistent with previous reports identifying c-Fos as a suppressor of hepatocyte differentiation [15, 26].

**Fig. 7.**
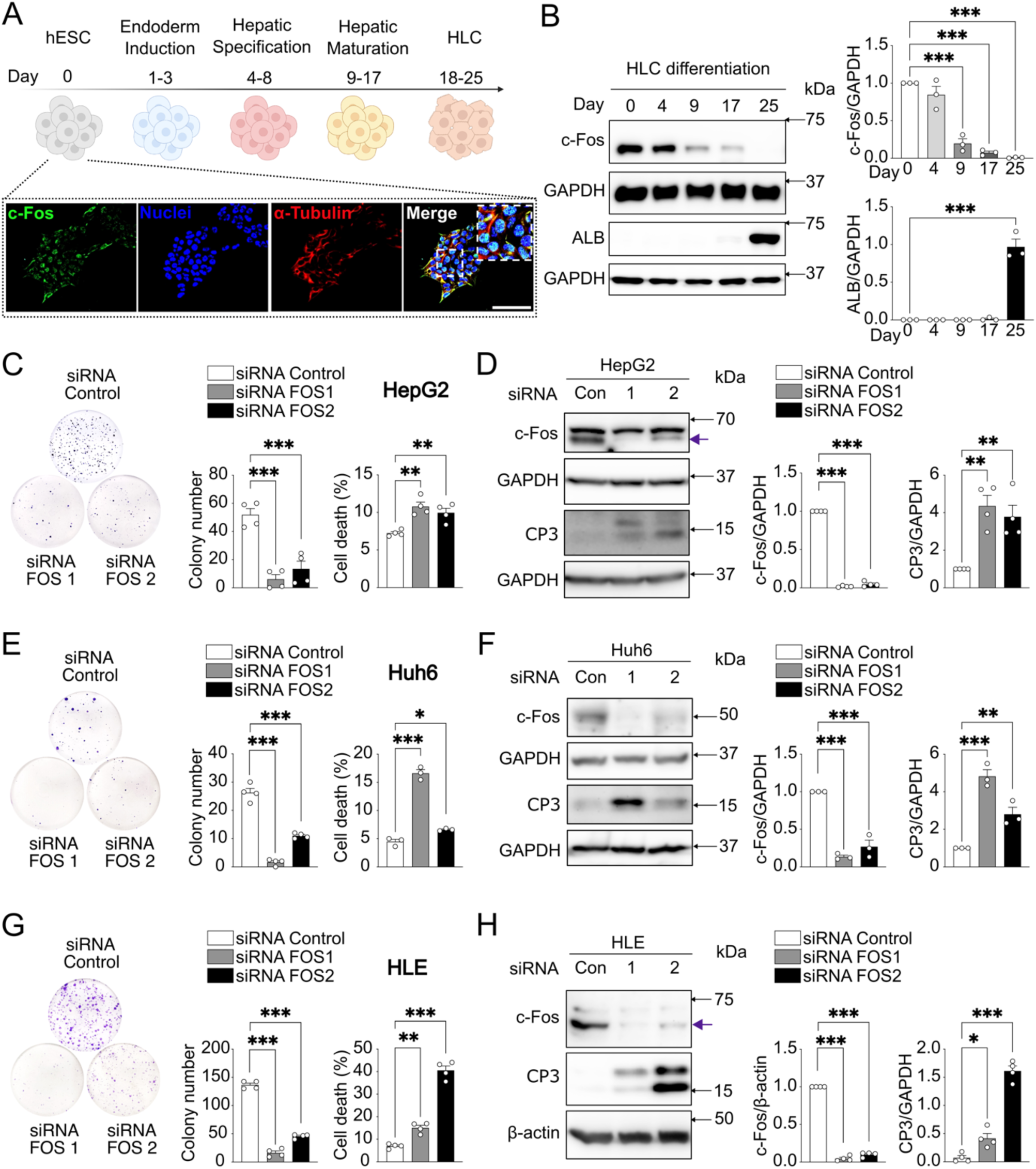
c-Fos expression reduces susceptibility of HCC cells to apoptosis. (A) Methodological approach schematic illustrating hepatocyte-like cells (HLC) differentiated from H1 human embryonic stem cells (hESC). Immunofluorescence showing c-Fos nuclear expression in stem cells. Scale bar=100μm. (B) Immunoblot analysis of HLC differentiation (n=3) showing c-Fos and hepatocyte marker ALB protein expression. (C, E, G) Colony formation capacity (left) and cell death (right) of HepG2 (C, n=4), Huh6 (E, n=3-4), and HLE (G, n=4) after siRNA-mediated c-Fos knockdown as indicated. (D, F, H) Immunoblot analysis of HepG2 (D, n=4), Huh6 (F, n=3), and HLE (H, n=4) showing c-Fos and cleaved caspase 3 (CP3) protein expression after siRNA-mediated c-Fos knockdown. Arrow in (D, H) indicating c-Fos. In (B-H), results are shown as mean ± SEM. Statistical analyses using one-way ANOVA (B-H). Statistical significance is indicated as \**p* < 0.05, \*\**p* < 0.01, \*\*\**p* < 0.001.

c-Fos is expressed across different human HCC cell lines (**Fig. S7A**). c-Fos siRNA silencing significantly reduced colony formation in all three cell lines tested, HepG2, Huh6, and HLE (**Fig. 7C**,**E**,**G**), accompanied by increased apoptosis, as evidenced by cleaved-caspase 3 induction (**Fig. 7D**,**F**,**H**). c-Fos has been implicated in promoting tumor invasion associated to JNK/c-Jun regulation [27–29]. Colony formation was inhibited at relatively high doses of the JNK inhibitor in HCC cell lines (**Fig. S7B-D**). These findings suggest that silencing c-Fos leads to decreased cell proliferation and increased apoptosis in these HCC cell lines, indicating c-Fos key role in contributing to the survival and proliferation of HCC cells.

### c-Fos protects HCC cells from apoptosis induced by lipotoxicity and ER stress

Saturated free fatty acids induce cell death in hepatocytes [30], but how cancer cells avoid lipotoxicity is unclear. To investigate whether c-Fos-mediated signaling counteracts lipotoxicity-induced apoptosis, we silenced c-Fos in HepG2 cells followed by palmitic acid treatment. This combination significantly increased cleaved caspase-3 (**Fig. 8A**), and cell death rates compared to palmitic acid treatment alone (**Fig. 8B**).

**Fig. 8.**
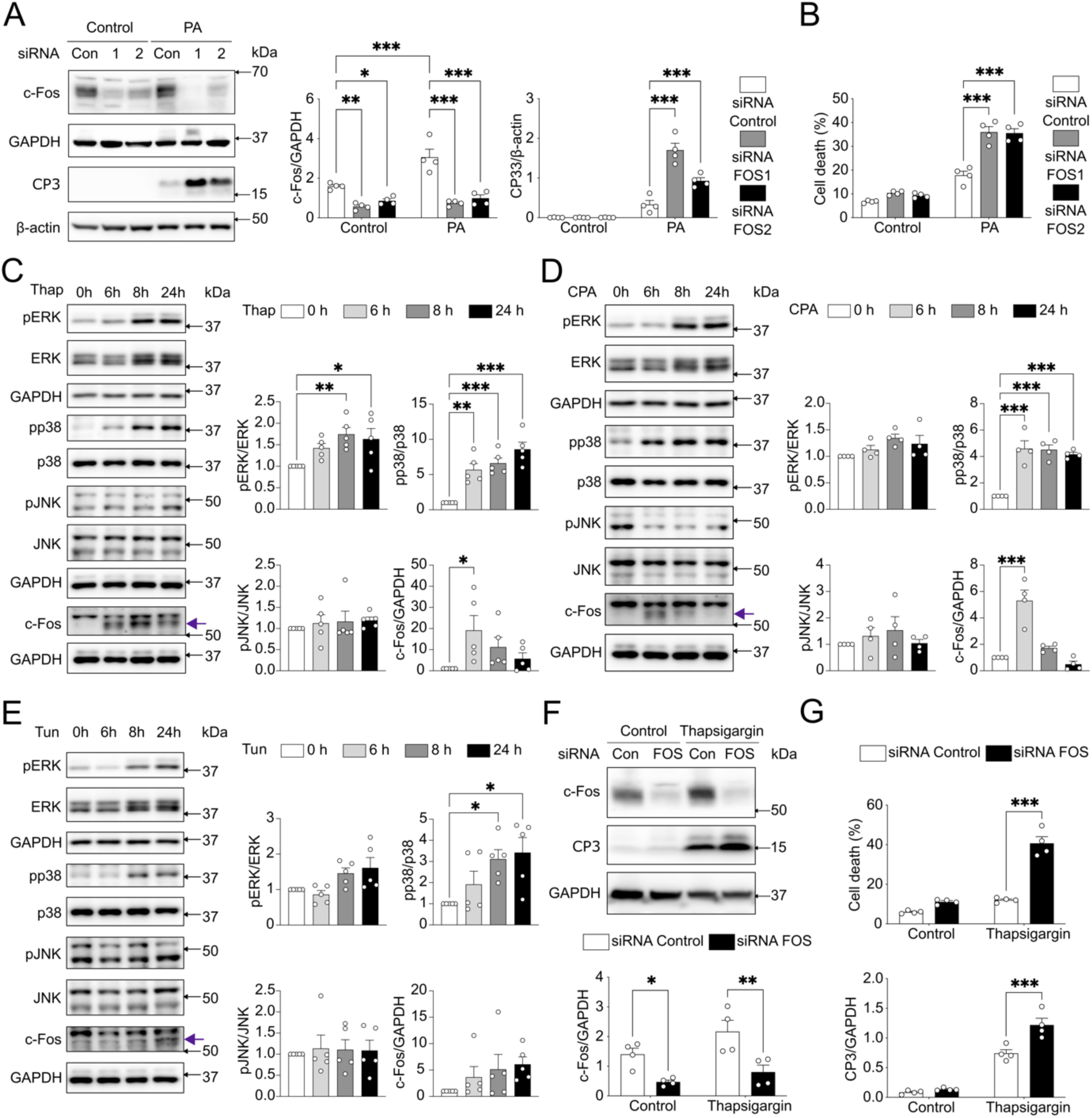
c-Fos knockdown induces ER stress susceptibility in HCC cells. (A, F) Immunoblot analysis of HepG2 treated with PA (A, 0.5mM, n=4) or thapsigargin (F, 1μM, n=4) after siRNA-mediated c-Fos knockdown, revealing c-Fos and cleaved caspase 3 (CP3) protein expression. (B, G) Cell death of HepG2 treated with PA (B, 0.5mM, n=4) or thapsigargin (G, 1μM, n=4) after siRNA-mediated c-Fos knockdown. (C-E) Immunoblot analysis of HepG2 treated with thapsigargin (Thap, C, n=5, 1μM), cyclopiazonic acid (CPA, D, n=4, 75μM), and tunicamycin (Tun, E, n=5, 5μg/mL), revealing pERK, pJNK, pp38 MAPK, and c-Fos protein expression. Arrow in (C-E) indicating c-Fos. In (A-G), results are shown as mean ± SEM. Statistical analyses using two-way (A, B, F, G) or one-way (C-E) ANOVA. Statistical significance is indicated as \**p* < 0.05, \*\**p* < 0.01, \*\*\**p* < 0.001.

Palmitic acid induces hepatic endoplasmic reticulum (ER) stress and disrupts Ca²⁺ signaling [30, 31]. HepG2 cells were treated with inhibitors (**Fig. S7E**) targeting Sarco/ER Ca²⁺-ATPase (SERCA) pump, including thapsigargin (**Fig. 8C**) and cyclopiazonic acid (**Fig. 8D**), as well as tunicamycin (**Fig. 8E**), an inhibitor of N-acetylglucosamine-1-phosphate transferase. c-Fos protein levels were transiently upregulated after 6h thapsigargin or cyclopiazonic acid treatment. However, p38 MAPK activation was consistently elevated across all treatments, while ERK activation was only observed with thapsigargin (**Fig. 8C**,**D**,**E**). Importantly, c-Fos siRNA silencing in HepG2 cells led to a marked increase in cell death after 24h of thapsigargin treatment (**Fig. 8F,G**). These results suggest that c-Fos activation protects from apoptosis in HCC cells exposed to lipotoxicity or SERCA-mediated ER stress.

We next analyzed liver RNA sequencing dataset generated using a mouse model, where c-Fos was specifically knocked out in hepatocytes during embryogenesis (Fos^Δli^) [15]. These mice are overall healthy, display histologically normal livers but are resistant to DEN-induced chemical carcinogenesis [15]. 8-week-old Fos^Δli^ and control mice on chow diet were injected with DEN and the livers were collected for RNA sequencing 48h later (**Fig. 9A**; **Fig. S8A**). At this early time point, liver function genes tended to be increased in Fos^Δli^ livers while carcinogenesis-related genes appeared decreased (**Fig. 9B**). Fos^Δli^/control GSEA revealed enriched KEGG pathways, including activated oxidative phosphorylation, suppressed pathways in cancer, cytokine-cytokine receptor interaction, chemokine signaling, calcium signaling, and focal adhesion (**Fig. 9C**). Consistent with previous results, these data suggest early protective effects of hepatic c-Fos deletion during carcinogenesis.

**Fig. 9.**
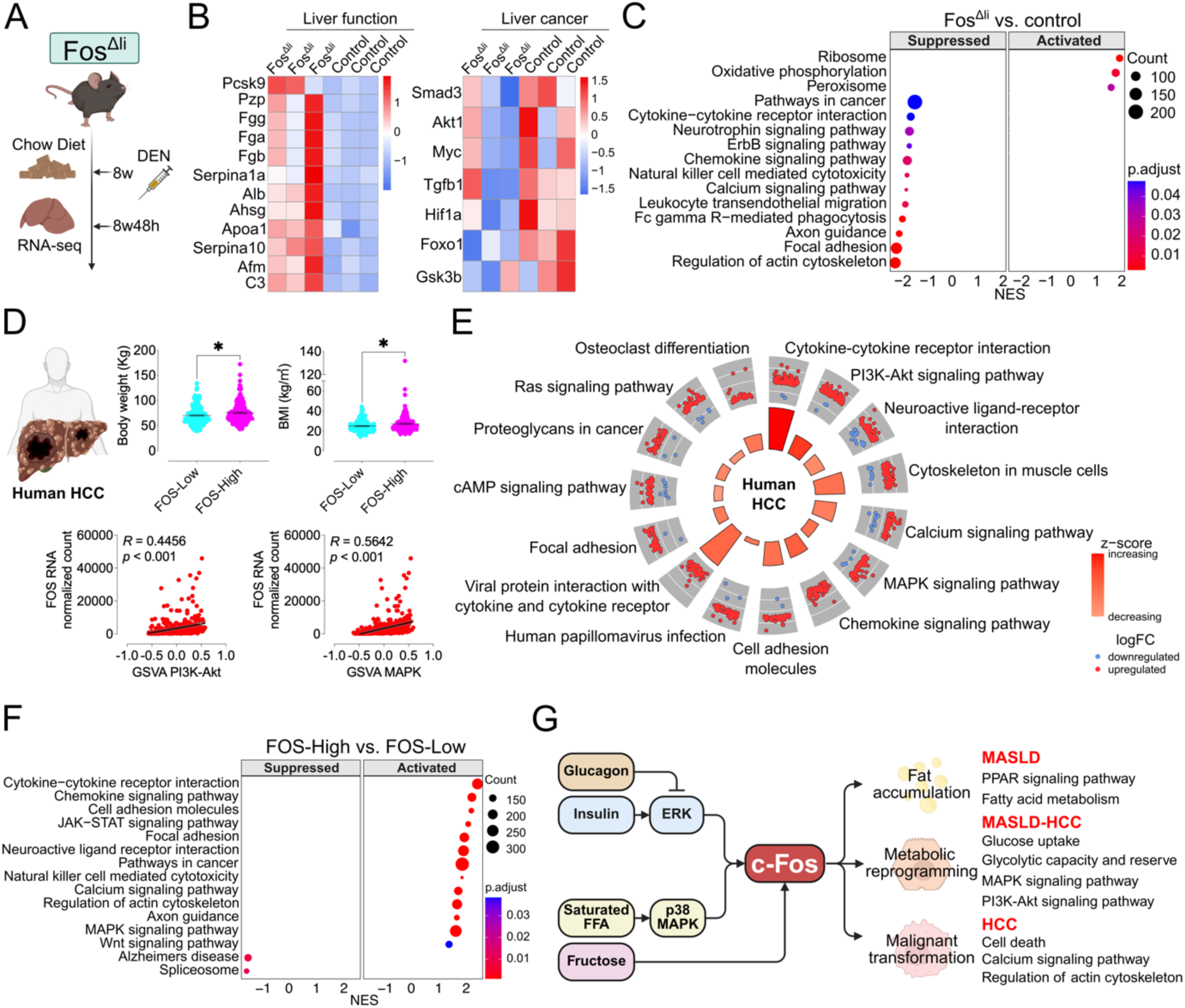
Mouse models with altered Fos expression hepatocytes reveal proximity to human MASLD and the development of HCC. (A) Hepatocyte-specific *Fos* knockout (Fos^Δli^) mice were subjected to RNA-Seq (GSE81079) after 8 weeks of chow diet and DEN injection for 48h. (B) RNA-Seq heatmap displaying liver function- and carcinogenesis-related gene alterations between Fos^Δli^ (n=3) and control (n=3). (C) RNA-Seq KEGG pathway enrichment analysis comparing Fos^Δli^ and control. (D) Human HCC data of TCGA was extracted (n=371). FOS-High (n=184) and -Low (n=187) groups were previously divided by median FOS expression counts in HCC RNA-Seq before using DESeq2 analysis. Body weight (FOS-High n=171, FOS-Low n=173) and BMI (FOS-High n=167, FOS-Low n=168) as indicated (top). FOS expression level showing a correlation with the customized PI3K-Akt and MAPK signaling GSVA scores in HCC RNA-Seq (n=371). (E) RNA-Seq circle plot showing the KEGG pathway enrichment comparing FOS-High (n=184) vs. FOS-Low (n=187). (F) RNA-Seq KEGG pathway enrichment analysis comparing FOS-High (n=184) vs. FOS-Low (n=187). (G) Schematic representation of the postulated c-Fos mechanism in hepatocytes, illustrating its role in modulating signaling pathways associated with MASLD to HCC. Differential expression analysis using DESeq2 and pathway enrichment analysis using fGSEA with Benjamini-Hochberg FDR correction (C, F). GSVA using Poisson kernel to estimate the expression distribution of each gene (D). Pathway enrichment analysis using clusterProfiler with Benjamini-Hochberg (E). Z-SCORE quantification using GOplot (E). In (D), results (top) are shown as mean ± SEM. Statistical analyses using two-tailed unpaired Student’s t-test (D). Correlation analyses using Spearman test (D). Statistical significance is indicated as \**p* < 0.05.

Finally, we investigated c-Fos-mediated hepatic signaling alterations in human HCC using RNA sequencing data from The Cancer Genome Atlas (TCGA) Liver Hepatocellular Carcinoma (LIHC) cohort obtained from UCSC Xena [32] and analyzed with DESeq2. Genes upregulated in PI3K-Akt and MAPK signaling pathways from Fos^Hep^ mice (described in **Fig. 6G**) were mapped to their human homologs and assessed using Gene Set Variation Analysis (GSVA) in the TCGA LIHC dataset. GSVA scores for the mouse-mapped PI3K-Akt and MAPK pathways positively correlated with FOS expression in human tumor tissues (*p* < 0.001) (**Fig. 9D**). Further analysis of human tumor tissues, incorporating clinical data, revealed that patients with relatively high FOS hepatic expression had also higher body weight (n=344) or higher body mass index (BMI) (n=335, 158/335 BMI > 25 kg/m²), consistent with increased FOS in obesity. ORA with z-score and GSEA identified several KEGG pathways, including cytokine-cytokine receptor interaction, calcium signaling, MAPK signaling, cell adhesion molecules, chemokine signaling, and focal adhesion, with upregulated genes in the FOS-High group all associated with malignant hepatocyte transformation (**Fig. 9E,F**). These data further support the validity of our findings using mouse samples and *in vitro* cell cultures to model human HCC (**Fig. 9G**).

## Discussion

In this study, we dissected the dynamic regulation of hepatic c-Fos by nutritional status and hormonal signals during fasting and feeding, revealing its critical role in regulating glucose and lipid metabolism under physiological and pathological conditions. We demonstrated that c-Fos influences lipid metabolism via key lipogenic markers such as PPARγ and PLIN2, both of which are central to lipid accumulation. Silencing c-Fos reduced hepatic fat accumulation, echoing the progression of metabolic dysfunction-associated liver diseases like MASLD and MASH. Our results are consistent with previous work, indicating that dysregulation of lipid metabolism in hepatocytes is intricately tied to transcriptional reprogramming driven by stress-responsive transcription factors [33, 34]. In this scenario, c-Fos plays a key metabolic role in the reprogramming of hepatocytes in obesity.

c-Fos is a key regulator of hepatocyte differentiation in organoid models, where Fos knockout promotes hepatocyte maturation [26]. We found that c-Fos expression is significantly elevated in the liver of obese mice, particularly during feeding, with its persistent activation linked to the p38 MAPK pathway. This suggests a metabolic shift in obesity, where stress-related pathways compensate for insulin resistance. These findings emphasize the role of c-Fos in driving metabolic dysfunction, as c-Fos upregulation correlates with increased lipogenesis and insulin resistance, evidenced by elevated levels of lipogenesis marker PPARγ and reduced glucose infusion rate during clamp. We have previously shown that JNK/c-Jun/BIM axis is a critical regulator of obesity-induced liver dysfunction [35]. Activation of JNK/c-Jun has been linked to hepatic steatosis, leading to increased BIM expression in hepatocytes, which contributes to mitochondrial dysfunction and insulin resistance [35]. Thus, c-Fos, another stress-responsive transcription factor of the AP-1 family, might function as a dimer with c-Jun in modulating liver metabolism and insulin signaling.

In HCC, c-Fos expression correlated with the activation of PI3K-Akt and MAPK pathways, two important mediators of tumor cell survival and metabolic reprogramming during HCC progression. Silencing c-Fos in HCC cell lines reduced cell proliferation and induced apoptosis, suggesting that c-Fos plays a crucial role in promoting cell survival under stress, particularly in lipotoxic and ER stress environments.

Our data are also in line with previous findings, showing that c-Fos plays a critical role in liver inflammation, hepatocyte proliferation, DNA damage response activation, and premalignant transformation [15]. We unraveled how c-Fos upregulation enhances glycolytic capacity and reserve; a metabolic shift commonly associated with malignant cells [36]. This suggests that c-Fos may drive premalignant transformation by promoting glycolysis, potentially through transcription-mediated metabolic reprogramming. Persistent c-Fos expression in hepatocytes contributes to preneoplastic transformation, as demonstrated using Fos^Hep^ and Fos^Δli^ mice [15], further emphasizing the dual role of c-Fos/AP-1 in metabolic regulation and liver carcinogenesis. In addition, c-Fos influences tumor metabolism by altering lipid biosynthesis and oxidative pathways in hepatic cancer models [5].

In conclusion, our study provides compelling evidence that c-Fos is a central regulator of metabolic pathways in the liver. c-Fos persistent activation under obesity and lipotoxicity promotes lipid accumulation, insulin resistance, and liver dysfunction.

### Limitations and Future Directions

Sex and age specific differences, unexplored here, could influence c-Fos dynamics, as estrogen is known to modulate lipid metabolism. Additionally, this study focuses on the role of c-Fos in hepatocytes, which may overlook the role of c-Fos in other liver cell types during MASLD-HCC development. Future studies should explore the development of specific inhibitors or gene-editing techniques to modulate c-Fos/AP-1 activity in liver cells, assessing their therapeutic potential in liver dysfunction and cancer progression.

## Materials and Methods

### Mice

The animal protocols were approved by the Commission of Ethics and Animal Welfare (CEBEA), Faculty of Medicine, Université libre de Bruxelles (Protocol No. 732, and 917). C57BL/6N mice were housed in an animal facility at the ambient temperature of 22 °C - 25 °C and 40% - 55% humidity, with the circadian of 12-hour light & 12-hour dark cycle and free access to diet and water, according to Belgian Regulations for Animal Care.

8-week-old male mice were assigned to different groups randomly with consideration of unbiased body weight. These mice were fed with either a chow low-fat diet (RM1 (P) 801151, Special Diets Services, UK) as a control or a high-fat, high-fructose, high-cholesterol diet (HFHFHCD, 40% kcal from fat, 20% kcal from fructose, 2% cholesterol; D09100310i, Research Diets, New Brunswick, NJ, USA).

### Body and liver composition measurement

The lean and fat mass of the mouse body and liver were measured based on nuclear magnetic resonance with the application of EchoMRI™ 3-in-1 body composition analyzer from EchoMedical Systems (Houston, TX, USA).

### DEN-induced HCC mouse model

2-week-old male mice were subjected to 1-time intraperitoneal injection into the lower abdomen with a 25 mg/kg DEN (dissolved in PBS) to establish a liver tumor model. The mice were maintained on a chow diet until 6 weeks old. Later, the mice were fed either a chow diet until 40 weeks old or an HFHFHCD until 31 weeks old, respectively. Tumor tissue and adjacent non-tumor tissue were collected after euthanizing the mice by cervical dislocation.

### Histological analysis

After euthanizing, the mouse liver tissues were collected, dissected, and rinsed with PBS. Then, the tissues were fixed in 4% PFA (pH 7.4), embedded in paraffin, and sectioned into 5-7µm slices by rotatory microtome, HistoCore MULTICUT R from Leica. After being placed on the positively charged slides, liver sections were stained with hematoxylin and eosin (H&E). Images were captured at 40x magnification by the NanoZoomer Digital Pathology system (Hamamatsu Photonics K.K., version SQ 1.0.9) for analysis.

### Primary mouse hepatocyte isolation, culture, treatment, and staining

Primary mHep were isolated from C57BL/6N mice with a two-step collagenase perfusion method through the vena cava. After opening the abdominal cavity, the infra-hepatic vena cava was cannulated for perfusion, and the portal vein was severed to drain the blood from the liver. In the first step, the liver was perfused with HBSS (without calcium and magnesium, Thermo Fisher Scientific, #14170138) supplemented with 10 mM HEPES–NaOH (pH 7.4) and saturated with O2/CO2 (95:5 vol/vol) at 37°C for 10 min. In the second step, collagenase type IV (0.3 mg/mL) was added to William’s E Medium (Thermo Fisher Scientific, #32551087), and perfusion continued for an additional 10 min, softening the liver tissue.

The softened liver was placed in a sterile dish, and the cells were dissociated using a coarse-toothed comb in cold William’s E Medium. The cell suspension was then passed through a 100-µm filter to remove cell clumps and centrifuged at 50g for 5 min at 4°C. The pellet was resuspended in William’s E Medium, layered onto a Percoll solution (Millipore Sigma, # GE17-0891-01), and centrifuged at 1000 RPM for 10 min.

The isolated primary mHep were seeded with attachment medium (William’s Medium with Glutamax, 10% FBS, 1% Penicillin-Streptomycin, and 10 mM HEPES). The 24-well plate was seeded with 100,000 cells in each well. The 8-well IBDID plate was seeded with 50,000 cells in each well. The 8-well Seahorse plate was seeded with 10,000 cells in each well. The attachment medium was changed to maintenance medium (William’s Medium with Glutamax, 10% FBS, 1% Penicillin-Streptomycin, 1% Non-Essential Amino Acids, 10 mM HEPES, and 5 µM Hydrocortisone) 3-4h after seeding and underwent overnight recovery.

Insulin (100 nM) and/or ERK inhibitor (150 nM) was added in mHep after 8h starvation media without serum (William’s Medium with Glutamax and 1% Penicillin-Streptomycin). Insulin (100 nM) and/or ERK inhibitor (150 nM) was added 8h after starvation without serum. The treatment with oleic acid (0.8 mM), palmitic acid (0.4 mM), and p38 MAPK inhibitor 1 (150 nM, Adezmapimod, HY-10256) or p38 MAPK inhibitor 2 (250 μM, Ergothioneine, HY-N1914) included 1% FBS and 1% fatty acid-free BSA in the media.

Immunofluorescent staining was done in an 8-well IBDID plate. After treatment, primary mHep were rinsed with PBS and fixed with 4% PFA for 15 min at room temperature. After blocking with UltraVision Protein Block (Thermo Scientific, # TA-060-PBQ), the primary antibody (**Supplementary Table S1**) was incubated with primary mHep at 4°C overnight or at room temperature for 2h. After rinsing the IBDID plate 3 times with PBS, a secondary antibody was added (**Supplementary Table S2**). After 1h of incubation at room temperature, the IBDID plate was rinsed with PBS 3 times again and then mounted with Antifade Mounting Medium with DAPI (VECTASHIELD^®^, # VEC.H-1200) for further observation under a fluorescent microscope, ZEISS AXIO Observer. D1.

### Adenovirus transduction

For in vitro experiments, primary mHep were seeded onto a 24-well plate for glucose uptake measurement and protein extraction, or an 8-well IBDID plate for lipid accumulation measurement. Adenovirus-mediated Adv-Control (Ad-CMV-Null, #1300, Vector Biolabs), Adv-c-Fos (Ad-CMV-m-FOS, SKU: ADV-259549, Vector Biolabs), and Adv-shRNA-Fos (Ad-GFP-U6-m-FOS-shRNA, #1835, Vector Biolabs) were transduced in primary mHep after overnight seeding. After 24h infection, the culture media was replaced by William’s Medium with 1.5 mM, 5.5 mM, 11.1 mM, or 22.2 mM glucose for glucose uptake measurement after 24h, or replaced by William’s Medium together with 1% BSA, 1% FBS, and 0.1 mM palmitic acid for lipid accumulation measurement after 24h. Glucose content in the medium was measured (GlucCell Glucose Monitoring System; Cesco Bioengineering). The primary mHep in IBDID plate were fixed with 4% PFA for 15 min at room temperature, stained with 5 μg/mL Nile Red solution for 20 min and then mounted with Antifade Mounting Medium with DAPI. Fluorescent images were measured using Fiji (Version: 2.14.0/1.54f) software, and the lipid accumulation area was calculated based on the mean area of each channel (Nile Red and DAPI). For in vivo experiments, chow diet-fed 6-week-old C57BL/6N male mice were infected with Adv-Control or Adv-shRNA-Fos by retro-orbital injection at a dose of 1.0 x 10^9^ PFU in 100 μl PBS.

### Extracellular acidification rates measurement during glycolytic stress test

Primary mHep were seeded in an 8-well Seahorse plate. After overnight recovery, the hepatocytes were transduced with Adv-c-Fos or Adv-shRNA-Fos for 24h in triplicate wells. Before the start of extracellular acidification rates (ECAR) measurement, the hepatocytes were equilibrated in XF DMEM Medium (5 mM HEPES, glucose- and pyruvate-free, #103575-100, Agilent Technologies) with 2 mM glutamine in a 37°C CO_2_-free incubator for 1h. Immediately before the ECAR measurement in the glycolytic stress test, the Seahorse plate was refreshed with XF DMEM Medium. During the glycolytic stress test, glucose (10 mM), oligomycin (10 μM), and 2-DG (in total 100 mM, divided into two consecutive injections of 50 mM) were dissolved in XF DMEM Medium and injected sequentially. The hepatocytes were collected immediately after the glycolytic stress test and stored at -80°C for protein extraction and immunoblot analysis. The ECAR measurements during the glycolytic stress test were conducted using the XFp Flux Analyzer (Seahorse Bioscience, Agilent Technologies).

### Hyperinsulinemic-euglycemic clamp

Mice were subjected to hyper-insulinemic/euglycemic clamp as described [37]. Briefly, 6-week-old male mice underwent jugular vein catheterization with exteriorization above the back using a single-channel vascular access magnetic button (Instech Laboratories, VABM1B/25), and then adenovirus was injected into the jugular vein through the established channel. After 12 days, the mice were subjected to clamp experiment. During clamp experiment, the insulin (Novo Nordisk, Actrapid^®^ 100 UI/ml) infusion was maintained at 1.5 mIU/kg body weight/min to induce hyperinsulinemia. Blood glucose was measured using the StatStrip® Xpress2™ glucometer (Nova Biomedical, #56506).

### RNA interference, colony formation, and cell death assay

HepG2, HLE, and Huh6 cell lines were cultured in DMEM (Thermo Fisher Scientific, #21885108) supplemented with 10% heat-inactivated FBS and 1% Penicillin-Streptomycin. For c-Fos knockdown, HCC cell lines were transfected with siRNAs targeting c-Fos or a negative control siRNA (working concentration: 30 nM; QIAGEN, Venlo, the Netherlands). The delivery of siRNA was after 24h of seeding, facilitated with Lipofectamine™ RNAiMAX Transfection Reagent (Thermo Fisher Scientific, #13778150) in Opti-MEM™ I Reduced Serum Medium (Thermo Fisher Scientific, #31985047) and DMEM with 10% FBS. The siRNA target sequences are provided in **Supplementary Table S3**. After 24h of siRNA transfection, HCC cells recovered for 24h before trypsinization to obtain a single-cell suspension. For the colony formation assays, 2,000 cells were seeded into P6 plates. Colonies were allowed to grow for 1-2 weeks, depending on the cell lines, and were fixed with 4% PFA and stained with 0.5% crystal violet. In the cell death assay, HCC cell lines recovered for 24h after siRNA transfection and were treated with/without palmitic acid (0.5 mM) for 24h. Cell death was assessed using SYTOX green (Thermo Fisher Scientific, Gibco, UK) at a final concentration of 5 μM.

### RNA extraction, RT-PCR, and transcriptomics

The mRNA from primary mHep was extracted with the Dynabeads™ mRNA DIRECT™ Purification Kit (Thermo Fisher Scientific, #61012). The total RNA from mouse liver tissues was extracted with the RNeasy Mini Kit (50) (QIAGEN, #74104) and the RNase-Free DNase Set (50) (QIAGEN, #79254). Reverse transcription from mRNA to cDNA was performed with the Reverse Transcription Core Kit (Eurogentec, #RT-RTCK-03). RT-PCR was conducted on the Bio-Rad CFX system (Bio-Rad Laboratories, Hercules, CA) with SsoAdvanced Universal SYBR^®^ Green Supermix (Bio-Rad, #1725274). Primer information is indicated in **Supplementary Table S4**. The liver total RNA was extracted as described above. The RNA integrity number (RIN) was measured using Agilent 5400. All the RINs were above 9.0. Total RNA quality analysis, library preparation (poly A enrichment), and sequencing were performed by Novogene (UK) company. NovaSeq X Plus Series (3Gb/sample, 10M PE reads of PE150) was used for sequencing.

### Western blot

RIPA Buffer (Cell Signaling Technology, #9806S) was used to extract total protein lysates from tissues. Cell Lysis Buffer (Cell Signaling Technology, #9803S) was used for extracting cell protein lysates. Both lysis buffers were supplemented with a Halt protease and phosphatase inhibitor cocktail (Thermo Fisher Scientific, #78442). Protein concentration was measured using a BCA protein assay kit (Thermo Fisher Scientific, #PI23227).

20 – 50 µg of protein lysate mixed with Laemmli buffer was loaded into the gel. The total amount of protein in each loading was the same in one gel. After electrophoretic separation in polyacrylamide gel, the protein was transferred onto a 0.22 µM nitrocellulose membrane (Bio-Rad, Hercules, CA, USA, #1620112). After blocking with 5% milk in 0.3% TBST, the primary antibody (detailed in **Supplementary Table S1**) was incubated with the membrane in 0.3% TBST with 5% BSA at 4°C overnight. A secondary antibody was added after washing the membrane 3 times with 0.3% TBST, including goat anti-rabbit IgG (Dako Agilent, #P0448), goat anti-mouse IgG (Dako Agilent, #P0447), and goat anti-guinea pig IgG (PROGEN Biotechnik, #90001) (detailed in **Supplementary Table S2**). For chemiluminescence detection, immunoreactive bands were visualized under the Amersham ImageQuant 800 Western blot imaging system (Cytiva Life Science, Marlborough, MA, USA).

### Bioinformatic analysis

The RNA sequencing reads of the mouse liver regarding GSE81079 were gathered in unaligned BAM format. The reads were converted to Fastq format by Picard and then aligned to the mouse reference genome (GRCm38.98). After the procedures of FastQC, fastp, Trimmomatic, HISAT2, SAMtools, and featureCounts, the reads were transformed into count matrix and underwent differential analysis by DESeq2 in R (4.4.1). The identified MASLD pathways, MASLD human proximity scores, and Fos expression data were extracted from PMCID: PMC11199145 source data. The human HCC dataset of the Cancer Genome Atlas Program (TCGA) was downloaded from UCSC Xena in aligned counts format and analyzed with differential analysis by DESeq2. KEGG pathway enrichment analysis of ORA (using clusterProfiler) with z-score (using GOplot), GSEA (using fGSEA), and customized pathway analysis of GSVA (using GSVA) followed the instructions on the corresponding websites. ORA was based on differentially expressed genes identified by DESeq2. GSEA was based on pre-ranked Wald statistics generated from DESeq2. ORA and GSEA were corrected with Benjamini-Hochberg. Poisson kernel was used to estimate the expression distribution in GSVA. The adjusted *p*-value < 0.05 was applied in differentially expressed genes and significantly enriched pathways.

### Stem cell differentiation in hepatocyte-like cells

Following a previously established protocol, stem cells were differentiated into hepatocyte-like cells [36]. Laminin 521-coated plates (BioLamina, Cat#LN521-05) were prepared, and human H1 embryonic stem cells were detached and seeded onto the laminin-coated plates (55,000 cells/well in P24). The cells were allowed to reach optimal confluency before initiating differentiation.

### Statistical analysis

The results are presented as mean ± standard error of the mean (SEM). Comparison between two groups was applied with two-tailed unpaired Student’s t-test. One-way ANOVA was employed for comparing more than two groups on a single independent variable. For comparisons between groups divided by two independent variables, two-way ANOVA was employed. Spearman test was used for correlation analyses between two parameters. Statistical analyses were conducted using Prism software (V.10, GraphPad Software, Inc, La Jolla, CA, USA). The sample size was predetermined based on variability observed in previous experiments and preliminary data. Statistical significance was defined as \**p* < 0.05, \*\**p* < 0.01, \*\*\**p* < 0.001.

## Acknowledgments

We thank Madalina Popa, André Dias, Anne Van Praet, Rabéa Dahili, Erick Arroba, Mariana Nunes, Francisco Costa, Cláudia S. Pinto, Tzu-Keng Shen, Leonardo Traini, Valerie Vandenbempt, Garnik Hovhannisyan, Carlos E. Buss, Javier Negueruela, Peng Xiao, and Xiaoyan Yi for experimental and technical support. We have used BioRender.com for the design of cartoons in the different panels (under license).

## Funding

The European Research Council (ERC) Consolidator grant METAPTPs grant Agreement No. GA817940 (ENG)

FNRS-WELBIO grant 35112672 (ENG)

FNRS-PDR grant 40007740 (ENG)

FNRS-TELEVIE grant 40007402 (ENG)

The ULB Foundation (ENG).

PhD scholarship support from China Scholarship Council (AL).

ENG is a Research Associate of the FNRS, Belgium.

## Author contributions

Conceptualization: AL, EHG, LB, ENG

Methodology: AL, EHG, WSW, CVD, SPS, RC

Visualization: AL, EHG, LH, LB, ENG

Supervision: EHG, ENG

Writing—original draft: AL, LH, EHG, ENG

Writing—review & editing: AL, LB, EHG, LH, ENG

## Data and materials availability

All data are available in the main text or the supplementary materials. Bulk RNA sequencing data can be downloaded from GEO (GSE297224). Source data regarding the reanalysis of RNA sequencing data can be found in the supplementary materials.

## References

1. Younossi, Z.M., et al., The global epidemiology of nonalcoholic fatty liver disease (NAFLD) and nonalcoholic steatohepatitis (NASH): a systematic review. Hepatology, 2023. 77(4): p. 1335–1347.

2. Talamantes, S., et al., Non-alcoholic fatty liver disease and diabetes mellitus as growing aetiologies of hepatocellular carcinoma. JHEP Rep, 2023. 5(9): p. 100811.

3. Brahma, M.K., et al., Oxidative stress in obesity-associated hepatocellular carcinoma: sources, signaling and therapeutic challenges. Oncogene, 2021. 40(33): p. 5155–5167.

4. Seyda Seydel, G., et al., Economic growth leads to increase of obesity and associated hepatocellular carcinoma in developing countries. Ann Hepatol, 2016. 15(5): p. 662–72.

5. Huang, R., et al., Multi-omics profiling reveals rhythmic liver function shaped by meal timing. Nat Commun, 2023. 14(1): p. 6086.

6. Bideyan, L., R. Nagari, and P. Tontonoz, Hepatic transcriptional responses to fasting and feeding. Genes Dev, 2021. 35(9-10): p. 635–657.

7. Beaulant, A., et al., Endoplasmic reticulum-mitochondria miscommunication is an early and causal trigger of hepatic insulin resistance and steatosis. J Hepatol, 2022. 77(3): p. 710–722.

8. Karin, M., Z. Liu, and E. Zandi, AP-1 function and regulation. Curr Opin Cell Biol, 1997. 9(2): p. 240–6.

9. Shaulian, E. and M. Karin, AP-1 as a regulator of cell life and death. Nat Cell Biol, 2002. 4(5): p. E131–6.

10. Curran, T., et al., FBJ murine osteosarcoma virus: identification and molecular cloning of biologically active proviral DNA. J Virol, 1982. 44(2): p. 674–82.

11. Wu, Q., et al., The temporal pattern of cfos activation in hypothalamic, cortical, and brainstem nuclei in response to fasting and refeeding in male mice. Endocrinology, 2014. 155(3): p. 840–53.

12. Madangopal, R., et al., Incubation of palatable food craving is associated with brain-wide neuronal activation in mice. Proc Natl Acad Sci U S A, 2022. 119(45): p. e2209382119.

13. Ray, J.D., et al., Nkx6.1-mediated insulin secretion and beta-cell proliferation is dependent on upregulation of c-Fos. FEBS Lett, 2016. 590(12): p. 1791–803.

14. Luther, J., et al., Fra-2/AP-1 controls adipocyte differentiation and survival by regulating PPARgamma and hypoxia. Cell Death Differ, 2014. 21(4): p. 655–64.

15. Bakiri, L., et al., Liver carcinogenesis by FOS-dependent inflammation and cholesterol dysregulation. J Exp Med, 2017. 214(5): p. 1387–1409.

16. Hasenfuss, S.C., et al., Regulation of steatohepatitis and PPARgamma signaling by distinct AP-1 dimers. Cell Metab, 2014. 19(1): p. 84–95.

17. Bu, L., et al., High-fat diet promotes liver tumorigenesis via palmitoylation and activation of AKT. Gut, 2024. 73(7): p. 1156–1168.

18. Vacca, M., et al., An unbiased ranking of murine dietary models based on their proximity to human metabolic dysfunction-associated steatotic liver disease (MASLD). Nat Metab, 2024. 6(6): p. 1178–1196.

19. Bakiri, L., et al., Functions of Fos phosphorylation in bone homeostasis, cytokine response and tumourigenesis. Oncogene, 2011. 30(13): p. 1506–17.

20. Mukherji, A., et al., An atlas of the human liver diurnal transcriptome and its perturbation by hepatitis C virus infection. Nat Commun, 2024. 15(1): p. 7486.

21. Keeton, A.B., et al., Insulin signal transduction pathways and insulin-induced gene expression. J Biol Chem, 2002. 277(50): p. 48565–73.

22. Messina, J.L., Insulin’s regulation of c-fos gene transcription in hepatoma cells. J Biol Chem, 1990. 265(20): p. 11700–5.

23. Mootha, V.K., et al., PGC-1alpha-responsive genes involved in oxidative phosphorylation are coordinately downregulated in human diabetes. Nat Genet, 2003. 34(3): p. 267–73.

24. Subramanian, A., et al., Gene set enrichment analysis: a knowledge-based approach for interpreting genome-wide expression profiles. Proc Natl Acad Sci U S A, 2005. 102(43): p. 15545–50.

25. Walter, W., F. Sánchez-Cabo, and M. Ricote, GOplot: an R package for visually combining expression data with functional analysis. Bioinformatics, 2015. 31(17): p. 2912–4.

26. Liang, J., et al., In-organoid single-cell CRISPR screening reveals determinants of hepatocyte differentiation and maturation. Genome Biol, 2023. 24(1): p. 251.

27. Eferl, R. and E.F. Wagner, AP-1: a double-edged sword in tumorigenesis. Nat Rev Cancer, 2003. 3(11): p. 859–68.

28. Hui, L., et al., Proliferation of human HCC cells and chemically induced mouse liver cancers requires JNK1-dependent p21 downregulation. J Clin Invest, 2008. 118(12): p. 3943–53.

29. Uhlirova, M. and D. Bohmann, JNK- and Fos-regulated Mmp1 expression cooperates with Ras to induce invasive tumors in Drosophila. Embo j, 2006. 25(22): p. 5294–304.

30. Zheng, S., et al., Calcium homeostasis and cancer: insights from endoplasmic reticulum-centered organelle communications. Trends Cell Biol, 2023. 33(4): p. 312–323.

31. Xu, W., et al., O-GlcNAc transferase promotes fatty liver-associated liver cancer through inducing palmitic acid and activating endoplasmic reticulum stress. J Hepatol, 2017. 67(2): p. 310–320.

32. Goldman, M.J., et al., Visualizing and interpreting cancer genomics data via the Xena platform. Nat Biotechnol, 2020. 38(6): p. 675–678.

33. Montagner, A., et al., Liver PPARalpha is crucial for whole-body fatty acid homeostasis and is protective against NAFLD. Gut, 2016. 65(7): p. 1202–14.

34. Blüher, M., Obesity: global epidemiology and pathogenesis. Nat Rev Endocrinol, 2019. 15(5): p. 288–298.

35. Litwak, S.A., et al., JNK Activation of BIM Promotes Hepatic Oxidative Stress, Steatosis, and Insulin Resistance in Obesity. Diabetes, 2017. 66(12): p. 2973–2986.

36. Gilglioni, E.H., et al., PTPRK regulates glycolysis and de novo lipogenesis to promote hepatocyte metabolic reprogramming in obesity. Nat Commun, 2024. 15(1): p. 9522.

37. Brenachot, X., et al., Hepatic protein tyrosine phosphatase receptor gamma links obesity-induced inflammation to insulin resistance. Nat Commun, 2017. 8(1): p. 1820.

