## Supplementary Figures and Tables for "Feeding induces c-Fos in hepatocytes contributing to hepatocellular carcinoma in obesity"

Ao Li *et al.*

**This PDF file includes:**

Figs. S1 to S8
Tables S1 to S4

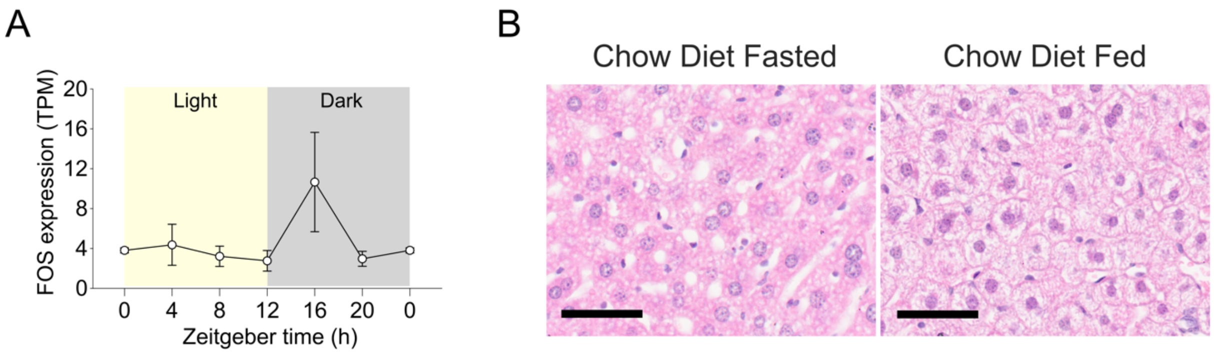


**Fig. S1. FOS expression and hepatocyte structure in fasted/fed livers.** (A) FOS expression fluctuates within 24h-cycle (zeitgeber time) in male humanized liver chimeric mice (PMCID: PMC11362569). Expression peaking within 4 h of the dark cycle start and returning to baseline within 8 h, reaching levels comparable to those observed during the light phase (n=3). (B) Representative H&E staining of mouse livers in fasted and fed groups with a chow diet as indicated. Scale bar=50um. In (A), results are shown as mean ± SEM.

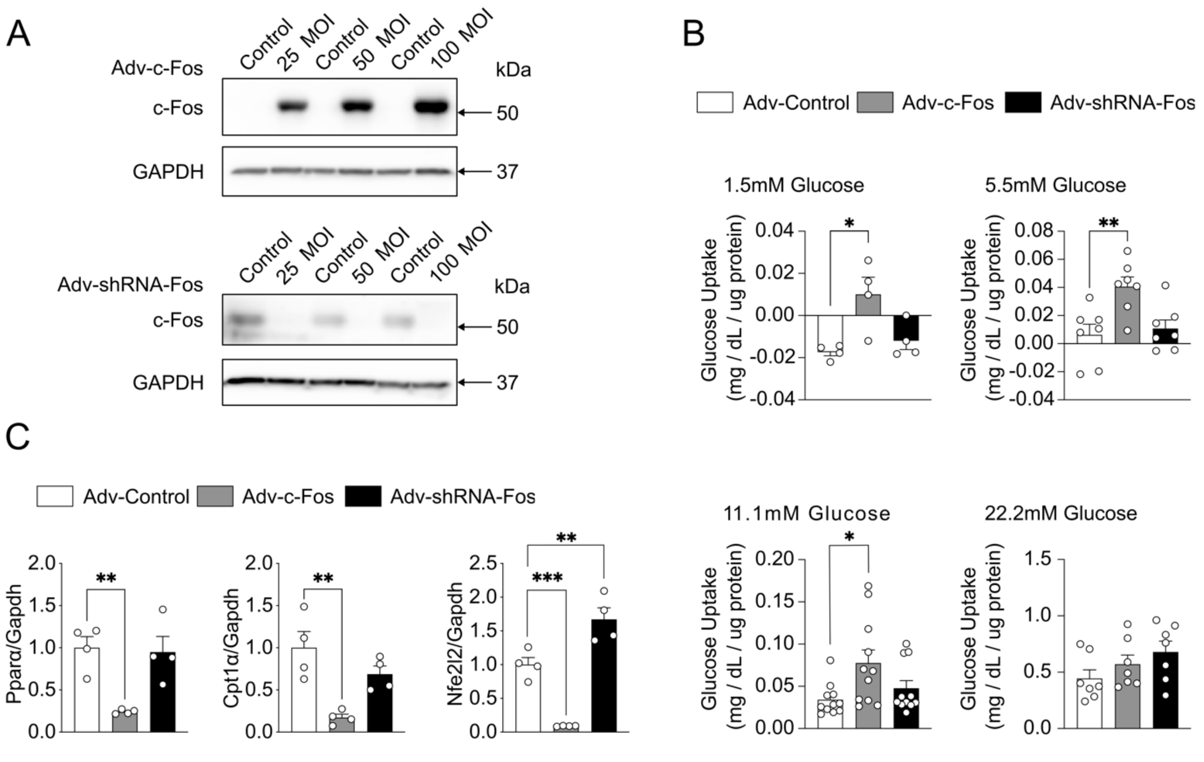


**Fig. S2. Adenovirus-mediated c-Fos overexpression enhances glucose uptake and suppresses β-oxidation-related genes expression in primary mouse hepatocytes.** (A) Representative immunoblot of primary mouse hepatocytes infected with adenoviral vectors to overexpress (Adv-c-Fos, top), silence (Adv-shRNA-Fos, bottom), or control (Adv-Control) for c-Fos expression modulation in different concentrations. 50 MOI was selected to overexpress or silence c-Fos in different experiments in this study. (B) Glucose uptake measurement in primary mouse hepatocytes upon overexpression and silencing of c-Fos as indicated (n=4-11). The primary mouse hepatocytes were cultured under 1.5mM, 5.5mM, 11.1mM, and 22.2mM of glucose Williams' E Medium for 24 h. (C) RT-PCR analysis revealing β-oxidation related genes *Pparα*, *Cpt1α*, and *Nfe2l2* expression in adenovirus-infected primary mouse hepatocytes with 22.2mM of glucose Williams' E Medium for 4 h (n=4). In (B, C), results are shown as mean ± SEM. Statistical analyses using one-way ANOVA (B, C). Statistical significance is indicated as *$p < 0.05$, **$p < 0.01$, ***$p < 0.001$.

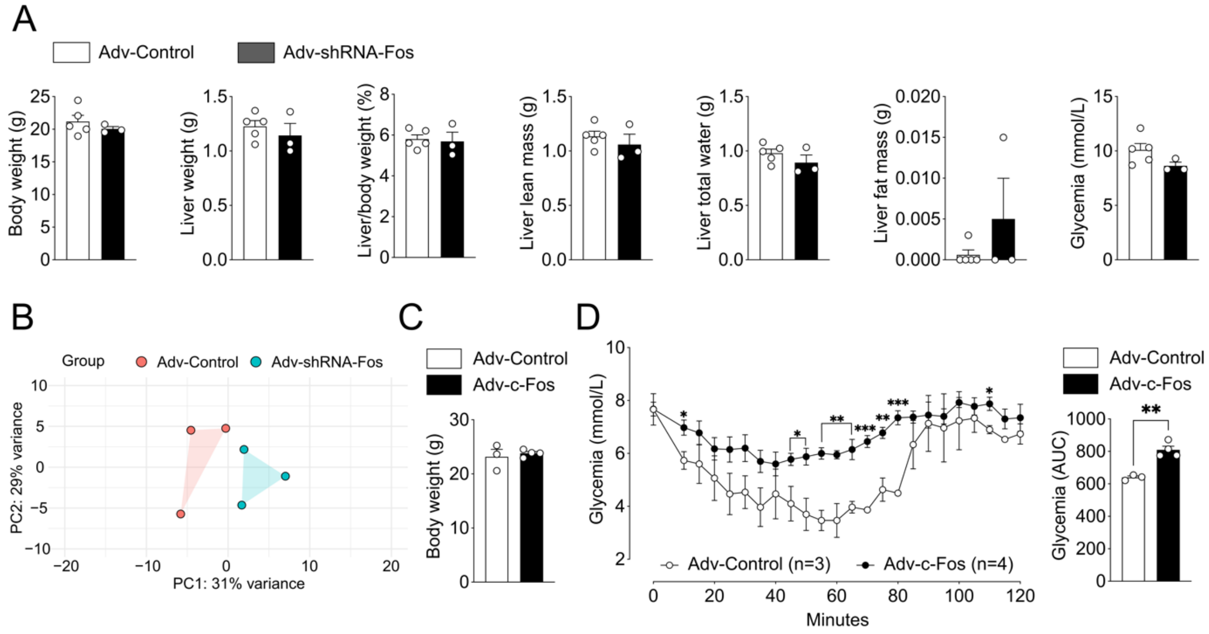


**Fig. S3. Short-term hepatic c-Fos manipulation alters liver genes expression and body insulin sensitivity in chow diet-fed mice.** (A) Metabolic parameters between Adv-Control (n=5) and Adv-shRNA-Fos (n=3) transduced mice. (B) Principal component analysis (PCA) plot using top 500 variable genes depicting the variation between Adv-Control (n=3) and Adv-shRNA-Fos (n=3) liver RNA-Seq data. PC1 and PC2 collectively explained 60% of the variability between Adv-shRNA-Fos and Adv-Control liver samples, with PC1 accounting for 31% and PC2 for 29% of the variance, respectively. (C) Body weight between Adv-Control (n=3) and Adv-c-Fos (n=4) mice in hyperinsulinemic/euglycemic clamp experiments as indicated. (D) Glycemia and area under curve (AUC) between Adv-Control (n=3) and Adv-c-Fos (n=4) during clamp as indicated. In (A, C, D), results are shown as mean ± SEM. Statistical analyses using two-tailed unpaired Student's t-test (A, C, D). Statistical significance is indicated as $^{**}p < 0.01$.

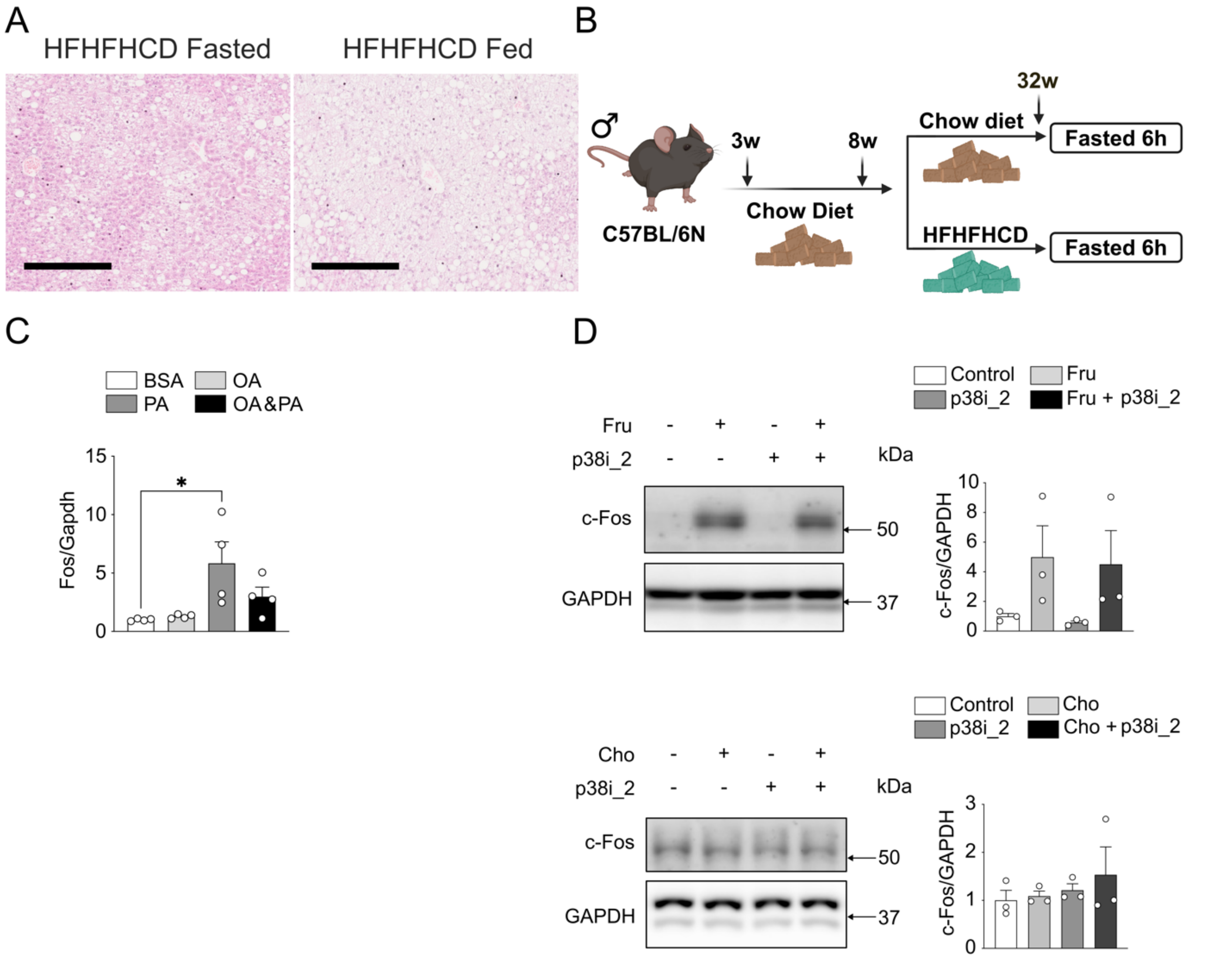


**Fig. S4. c-Fos is induced by saturated fatty acid palmitic acid and fructose in hepatocytes.** (A) Representative H&E staining of mouse livers in fasted and fed groups with HFHFHCD diet as indicated. Scale bar=250um. (B) Methodological approach schematic illustrating the mice fed with 25-week HFHFHCD or a chow diet starting from 8-week to 32-week. (C) RT-PCR analysis shows *Fos* mRNA expression after 24 h of palmitic acid (PA, 0.4 mM), oleic acid (OA, 0.8 mM), or PA&OA treatment in primary mouse hepatocytes (n=4). (D) Immunoblot analysis of primary mouse hepatocytes treated with fructose (Fru, 22.2 mM, n=3, top) or cholesterol (Cho, 0.1 µM, n=3, bottom) with p38i_2 (250 µM). In (C, D), results are shown as mean ± SEM. Statistical analyses using one-way ANOVA (C, D). Statistical significance is indicated as *$p < 0.05$.

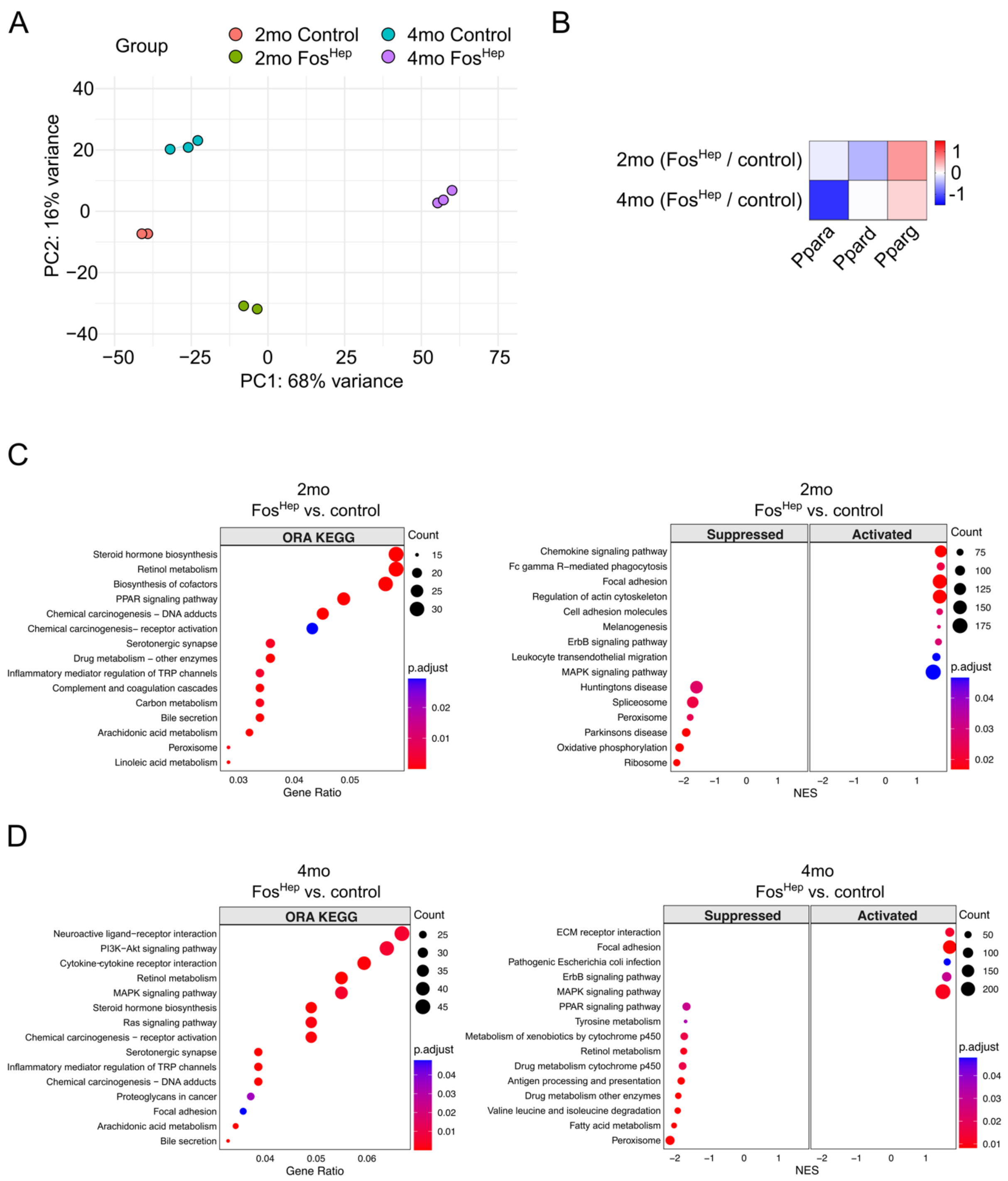


**Fig. S5. Hepatic c-Fos expression significantly alters KEGG signaling pathways.** (A) RNA-Seq PCA plot using top 5000 variable genes depicting the variation among 2 months (2mo, control n=2, $Fos^{Hep}$ n=2) and 4 months (4mo, control n=3, $Fos^{Hep}$ n=3). PC1 and PC2 collectively explained 84% of the variability among different groups, with PC1 accounting for 68% and PC2 for 16% of the variance, respectively. (B) PPAR genes expression in $Fos^{Hep}$/control mouse livers in 2mo and 4mo as indicated with $log_2$ fold change. (C, D) RNA-Seq KEGG pathway enrichment analysis comparing $Fos^{Hep}$ vs. control in 2mo (C) and 4mo (D), respectively. KEGG top 15 pathways ranked by gene count in corresponding pathways (ORA left, GSEA right). Differential expression analysis using DESeq2 and pathway enrichment analysis using clusterProfiler (left of C and D) and fGSEA (right of C and D) with Benjamini-Hochberg.

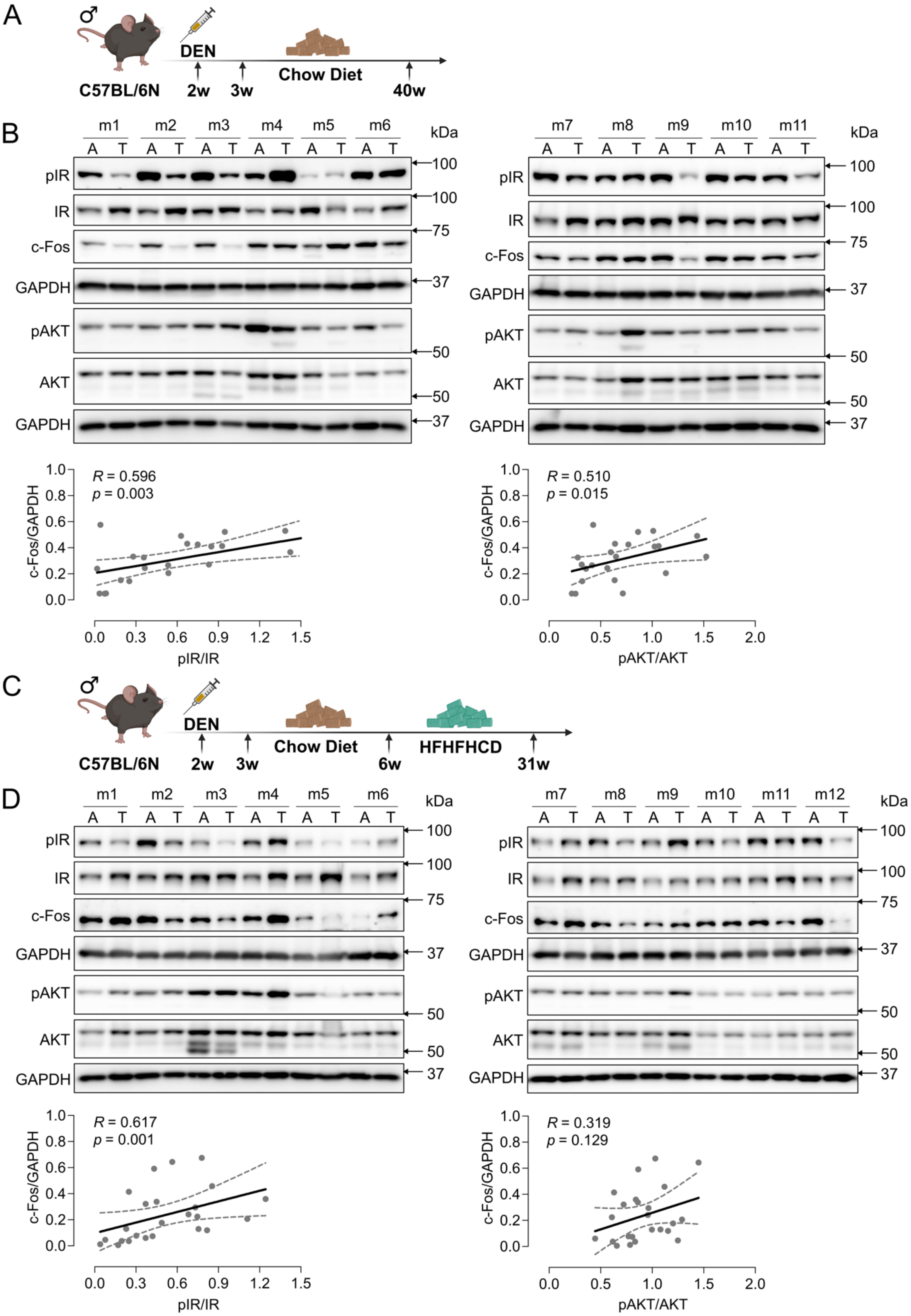
A
♂
C57BL/6N
DEN
2w
3w
Chow Diet
40w
B
m1 m2 m3 m4 m5 m6
A T A T A T A T A T A T
kDa
pIR
IR
c-Fos
GAPDH
pAKT
AKT
GAPDH
100
100
75
37
50
50
37
m7 m8 m9 m10 m11
A T A T A T A T A T
R = 0.596
p = 0.003
c-Fos/GAPDH
pIR/IR
R = 0.510
p = 0.015
pAKT/AKT
C
♂
C57BL/6N
DEN
2w
3w
Chow Diet
6w
HFHFHCD
31w
D
m1 m2 m3 m4 m5 m6
m7 m8 m9 m10 m11 m12
A T A T A T A T A T A T
kDa
pIR
IR
c-Fos
GAPDH
pAKT
AKT
GAPDH
R = 0.617
p = 0.001
c-Fos/GAPDH
pIR/IR
R = 0.319
p = 0.129
pAKT/AKT

**Fig. S6. Hepatic c-Fos expression correlates with phosphorylation of insulin receptor in DEN-induced HCC male mouse models.** (A, C) Methodological approach schematic illustrating diethyl nitrosamine (DEN)-induced HCC male mouse models fed with 37-week chow diet (A, 3-40 weeks duration) or 25-week HFHFHCD (C, 6-31 weeks duration) in this study. (B, D) Liver adjacent (labelled as A above immunoblot) and tumor tissues (labelled as T above immunoblot) were collected from chow diet (B, n=11) or HFHFHCD (D, n=12) fed DEN-induced HCC mice at end points, respectively. Immunoblot analysis of liver adjacent and tumor tissues showing the protein expression levels of pIR, pAKT, and c-Fos. Correlation analyses using Spearman test (B, D).

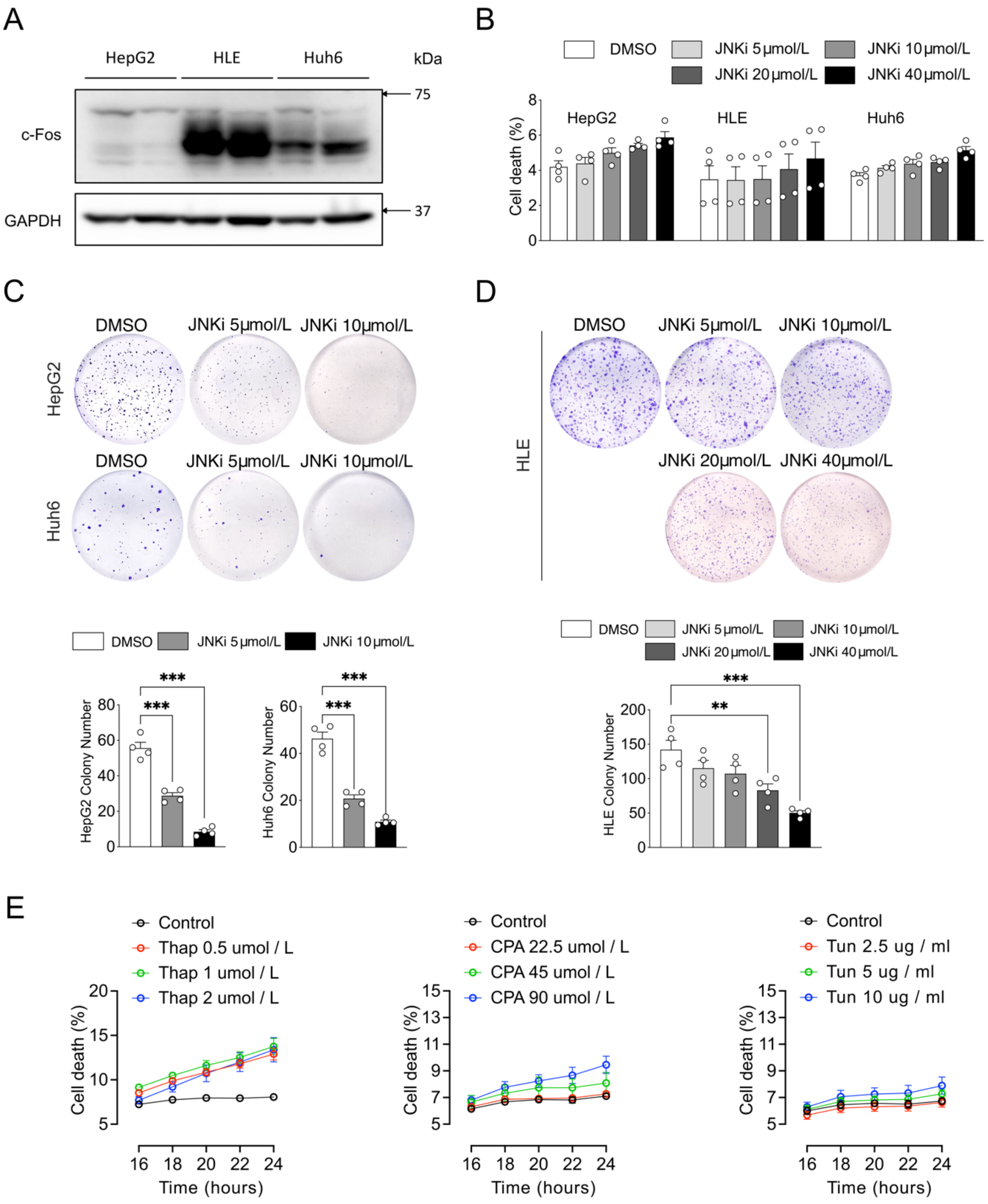


**Fig. S7. c-Fos expression in HCC cell lines, JNK inhibitor-mediated cell death and colony formation, and ER stressors induced cell death.** (A) Immunoblot analysis of HepG2, Huh6, and HLE HCC cell lines showing c-Fos protein expression levels. (B) Cell death assay showing different dosages of JNK inhibitor (JNKi) treatment on viability capacity of HepG2, Huh6, and HLE as indicated (n=4 for each cell line). (C, D) Colony formation assay of HepG2 (C), Huh6 (C), and HLE (D) treated with different dosages of JNKi as indicated (n=4 for each cell line). (E) Cell viability assay of HepG2 treated with different dosages of ER stressors, including Thap, Tun, and CPA, as indicated (n=4 for each cell line). In (B-E), results are shown as mean ± SEM. Statistical analyses using one-way ANOVA (B-D). Statistical significance is indicated as $^{**}p < 0.01$, $^{***}p < 0.001$.

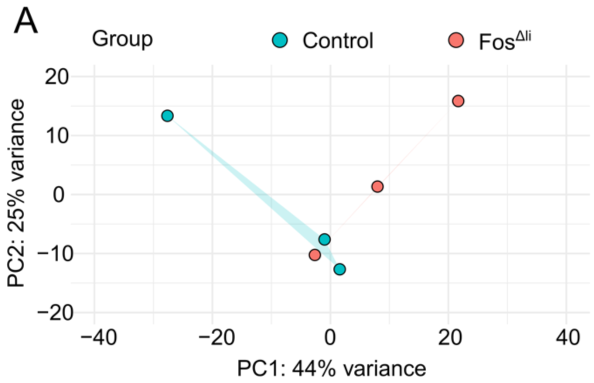


**Fig. S8. RNA-Seq PCA plot reveals variation between control and Fos$^{\Delta li}$ mice.** (A) RNA-Seq PCA plot using top 5000 variable genes depicting the variation between control and Fos$^{\Delta li}$. PC1 and PC2 collectively explained 69% of the variability among different groups, with PC1 accounting for 44% and PC2 for 25% of the variance, respectively.

| Primary antibody | Designation | Source or reference | Cat# | Additional information |
|---|---|---|---|---|
| AKT | Akt (pan) (40D4) Mouse mAb | Cell Signaling Technology | 2920 | WB 1:2000 |
| ALB | Albumin Antibody | Cell Signaling Technology | 4929 | WB 1:1000 |
| c-Fos | Rabbit polyclonal anti-protein c-Fos | Cell Signaling Technology | 4384S | WB 1:1000 |
| c-Fos | Rabbit monoclonal anti-protein c-Fos | Cell Signaling Technology | 2250S | WB 1:1000 IF 1:125 |
| Cleaved Caspase 3 | Rabbit monoclonal anti-protein cleaved caspase 3 | Cell Signaling Technology | 9664S | WB 1:1000 |
| ERK | p44/42 MAPK (Erk1/2) Antibody | Cell Signaling Technology | 9102 | WB 1:1000 |
| GAPDH | Rabbit polyclonal anti-glyceraldehyde-3-phoshphate dehydrogenase | R&D Systems | 2275-PC-100 | WB 1:5000 |
| IR | Insulin Receptor β (4B8) Rabbit mAb | Cell Signaling Technology | 3025 | WB 1:1000 |
| JNK | SAPK/JNK Antibody | Cell Signaling Technology | 9252 | WB 1:1000 |
| p-AKT | Phospho-Akt (Ser473) (D9E) XP® Rabbit mAb | Cell Signaling Technology | 4060 | WB 1:2000 |
| p-eIF2α | Phospho-eIF2α (Ser51) (119A11) Rabbit mAb | Cell Signaling Technology | 3597 | WB 1:1000 |
| p-ERK | Phospho-p44/42 MAPK (Erk1/2) (Thr202/Tyr204) (D13.14.4E) XP® Rabbit mAb | Cell Signaling Technology | 4370 | WB 1:2000 |
| p-IR | Phospho-IGF-I Receptor β (Tyr1135/1136)/Insulin Receptor β (Tyr1150/1151) (19H7) Rabbit mAb | Cell Signaling Technology | 3024 | WB 1:1000 |
| p-JNK | Phospho-SAPK/JNK (Thr183/Tyr185) Antibody | Cell Signaling Technology | 9251 | WB 1:1000 |
| p-p38 | Phospho-p38 MAPK (Thr180/Tyr182) (D3F9) XP® Rabbit mAb | Cell Signaling Technology | 4511 | WB 1:1000 |
| p38 | p38 MAPK (D13E1) XP® Rabbit mAb | Cell Signaling Technology | 8690 | WB 1:1000 |
| PLIN2 | Guinea pig polyclonal anti-protein Perilipin 2 (N-terminus aa 6-27) | PROGEN Biotechnik | GP47 | WB 1:1000 |
| PPARγ | Rabbit monoclonal anti-PPARγ | Cell Signaling Technology | 2443 | WB 1:500 |
| α-Tubulin | Mouse monoclonal anti-alpha-tubulin | Sigma-Aldrich | T5168 | IF 1:500 |
| β-actin | Anti-β-Actin antibody, Mouse monoclonal | Sigma-Aldrich | A1978 | WB 1:5000 |

**Table S1.** List of primary antibodies used for Western blot and immunofluorescence analysis.

| Secondary antibody | Designation | Source or reference | Cat# | Additional information |
|---|---|---|---|---|
| Donkey anti-Mouse 555 | Donkey anti-Mouse IgG (H+L) Highly Cross-Adsorbed Secondary Antibody, Alexa Fluor Plus 555 | Thermo Fisher Scientific | A32773 | IF 1:500 |
| Donkey anti-Rabbit 488 | Donkey anti-Rabbit IgG (H+L) Highly Cross-Adsorbed Secondary Antibody, Alexa Fluor 488 | Thermo Fisher Scientific | A21206 | IF 1:500 |
| Goat anti-guinea pig | Anti-guinea pig IgG goat polyclonal, HRP conjugate | PROGEN Biotechnik | 90001 | WB 1:5000 |
| Goat anti-Mouse | Goat Anti-Mouse Immunoglobulins/HRP (affinity isolated) | Dako | P0447 | WB 1:5000 |
| Goat anti-Rabbit | Goat Anti-Rabbit Immunoglobulins/HRP (affinity isolated) | Dako | P0448 | WB 1:5000 |

**Table S2.** List of secondary antibodies used for Western blot and immunofluorescence analysis.

| siRNA | Source | Identifiers | Target sequence |
|---|---|---|---|
| siRNA Control | Qiagen | #1027281 | - |
| Hs siRNA FOS #1 | Qiagen | SI03066028 | CAGCATGGAGCTGAAGACCGA |
| Hs siRNA FOS #2 | Qiagen | SI03091844 | CTCGGGCTTCAACGCAGACTA |

**Table S3.** siRNAs used in the study to knock down c-Fos in HCC cell lines.

| Gene | qPCR F | qPCR R | Standard F | Standard R |
|---|---|---|---|---|
| Acaca | AGCCAGAAGGGACAGTAGAA | CTCAGCCAAGCGGATGTAAA | GCGCTTACATTGTGGATGGC | AAGCCTTCACTGTGCCTTCA |
| Acly | TTCGTCAAACAGCACTTCC | ATTTGGCTTCTTGGAGGTG | ACACCATCATCTGTGCTCGG | ATCCCAGGGGTGACGATACA |
| Cpt1α | GCTGATGACGGCTATGGTGT | AAAGCGGTGTGAGTCTGTCT | CCACAACAACGGCAGAGCA | TCAGGAGCAACACCTATTCATTTG |
| Cyp7a1 | TAAGGAGAAGGAAAGTAGGTGAAC | CCAAATGCCTTCGCAGAAGTAG | TGGAATAAGGAGAAGGAAAGTAGG | TCCAAATGCCTTCGCAGAAGTA |
| Fasn | CACAGTGCTCAAAGGACATGCC | CACCAGGTGTAGTGCCTTCCTC | ACTTCCTCTGGGATGTGCCT | GTCAGCACTGCTCTCGTTGA |
| Fos | AGCAGCTATCTCCTGAAGAGG | TCTGTCTCCGCTTGGAGTGT | CTCCTGTCAACACACAGGACTT | GTTGATCTGTCTCCGCTTGGA |
| Gapdh | AGTTCAACGGCACAGTCAAG | TACTCAGCACCAGCATCACC | ATGACTCTACCCACGGCAAG | TGTGAGGGAGATGCTCAGTG |
| Nfe212 | ACTACAGTCCCAGCAGAGTGAT | TCACACACTTTCTGCGTGCT | ACTACAGTCCCAGCAGAGTGATG | AGACACTGCACTGCAACAAG |
| Nr1h3 | TTCTCCTGATTCTGCAACGGA | GACGAAGCTCTGTCGGCTCT | TTCTCCTGATTCTGCAACGGA | GCTTTTGTGGACGAAGCTCTG |
| Pparα | GCTGTAAGGGCTTCTTTCGG | GCGAATTGCATTGTGTGACAT | GCATGTGAAGGCTGTAAGGGC | GACAAAAGGCGGGTTGTTGCT |
| Pparγ1 | CCAAGAATACCAAAGTGCGATCA | AAAACCCTTGCATCCTTCACAA | GCTCCAAGAATACCAAAGTGCGA | AACCTGATGGCATTGTGAGACA |
| Pparγ2 | TGCCTATGAGCACTTCACAAG | TCTACTTTGATCGCACTTTGGTA | AGCATGGTGCCTTCGCTGAT | GCCCAAACCTGATGGCATTGTG |

**Table S4.** List of primers used for qPCR**.** Real-time quantitative PCR was performed using the Bio-Rad CFX96 machine (Bio-Rad Laboratories, Hercules, CA) and the SYBR green PCR Master Mix (Bio-Rad Laboratories).
